# Surfactin stimulated by pectin molecular patterns and root exudates acts as a key driver of *Bacillus*-plant mutualistic interaction

**DOI:** 10.1101/2021.02.22.432335

**Authors:** Grégory Hoff, Anthony Arguelles-Arias, Farah Boubsi, Jelena Prsic, Thibault Meyer, Heba M. M. Ibrahim, Sebastien Steels, Patricio Luzuriaga, Aurélien Legras, Laurent Franzil, Michelle Lequart, Catherine Rayon, Victoria Osorio, Edwin de Pauw, Yannick Lara, Estelle Deboever, Barbara de Coninck, Philippe Jacques, Magali Deleu, Emmanuel Petit, Olivier Van Wuytswinkel, Marc Ongena

**Author notes:** Grégory Hoff, Marc Ongena.

## Abstract

*Bacillus velezensis* is considered as model species belonging to the so-called *B. subtilis* complex that typically evolved to dwell in the soil rhizosphere niche and establish intimate association with plant roots. This bacterium provides protection to its natural host against diseases and represents one of the most promising biocontrol agents. However, the molecular basis of the cross-talk that this bacterium establishes with its natural host has been poorly investigated. We show here that these plant-associated bacteria have evolved some polymer-sensing system to perceive their host and that in response, they increase the production of the surfactin-type lipopeptide. Furthermore, we demonstrate that surfactin synthesis is favoured upon growth on root exudates and that this lipopeptide is a key component used by the bacterium to optimize biofilm formation, motility and early root colonization. In this specific nutritional context, the bacterium also modulates qualitatively the pattern of surfactin homologues co-produced *in planta* and mainly forms variants that are the most active at triggering plant immunity. Surfactin represents a shared good as it reinforces the defensive capacity of the host.

**Importance:** Within the plant-associated microbiome, some bacterial species are of particular interest due to the disease protective effect they provide via direct pathogen suppression and/or stimulation of host immunity. While these biocontrol mechanisms are quite well characterized, we still poorly understand the molecular basis of the cross talk these beneficial bacteria initiate with their host. Here we show that the model species *Bacillus velezensis* stimulates production of the surfactin lipopeptide upon sensing pectin as cell surface molecular pattern and upon feeding on root exudates. Surfactin favors bacterial rhizosphere fitness on one hand and primes the plant immune system on the other hand. Our data therefore illustrate how both partners use this multifunctional compound as unique shared good to sustain mutualistic interaction.

## Introduction

Soil is among the richest ecosystems in terms of microbial diversity, but only a subset of these microbes has evolved to efficiently establish in the competitive and nutrient-enriched rhizosphere layer surrounding plant roots (1). The rhizosphere includes plant beneficial bacteria dwelling on the rhizoplane as multicellular biofilm communities, feeding on exuded carbohydrates (2, 3), and, in turn, contributing to host fitness via growth stimulation and protection against phytopathogens (4, 5). This biocontrol activity is mediated via competition for nutrients and space, direct growth inhibition of the pathogenic (micro)organisms and more indirectly, by stimulating the host defensive capacity in an immunization-like process which leads to induced systemic resistance (ISR, (6, 7)). This ISR mechanism results in enhanced defense lines and reduced disease symptoms upon perception of plant beneficial microbes (6, 8).

From an ecological viewpoint, rhizosphere establishment and persistence of these beneficial bacteria rely on various traits but efficient root colonization and high competitiveness toward the surrounding microbiological network are pivotal. It is hypothesized that the potential to produce a wide range of chemically diverse and bioactive secondary metabolites (BSMs) acting as signals and/or antimicrobials is a common key feature of these beneficial bacteria (5, 9, 10). Members of the *Bacillus velezensis* species are considered as archetypes of plant-associated beneficial bacilli and are among the most prolific BSMs producers with more than 12% of their genome devoted to the synthesis of compounds contributing to both ecological competence and biocontrol activity (11–15). Among their BSM arsenal, the cyclic lipopeptide surfactin, is synthesized non-ribosomally by a multi-modular mega-enzyme machinery (encoded by the *srfA* operon) and is formed as a mix of naturally co-produced homologues varying in the length of the fatty acid chain. This multifunctional compound is of particular interest because it retains important roles in key developmental processes such as bacterial motility, biofilm formation and root colonization (16–18), but also because it represents the best described *Bacillus* triggers for plant immunity (6, 8). The potential of surfactin to stimulate ISR has been demonstrated on various plants including Solanaceae like tobacco and tomato on which it acts as main if not sole elicitor formed by *B. subtilis* and *B. velezensis* species (10, 19). In support to its key role in interaction with the host plant, we also previously reported that surfactin is promptly formed in the course of early colonization and that its production is stimulated upon sensing root tissues (20).

However, in contrast to the well-studied interactions between plants and microbial pathogens or nitrogen-fixing bacteria (21), relatively little is known on the molecular basis of cooperative interactions between plants and beneficial bacteria such as *B. velezensis* (11, 20, 22). More specifically, how and to what extent the expression of key bacterial BSMs may be modulated by plant factors is poorly understood. A better knowledge is not only critical for providing new insights in rhizosphere chemical ecology but also for optimizing the use of these species as biocontrol agents, which still suffer from insufficient efficacy in practice (23). Here, we investigated the molecular interaction driving the early steps of partnership establishment between plant roots and *B. velezensis*. We show that cell wall pectin acts in synergy with soluble root exudates as plant host cues perceived by *B. velezensis*. In response, the bacterium stimulates the production of specific surfactin variants as key components of its secretome to further improve the fitness of both partners *i*.*e*. early root colonization and thus rhizosphere competence of the bacterium and priming of immunity in the host plant.

## Results

### Pectin fragments of high polymerization degree act as host cues triggering surfactin production

We previously described that early production of surfactin, as a mix of naturally co-produced homologues varying in the length of the fatty acid chain, is stimulated in contact with root tissues and several plant cell wall-associated polymers (PCWP) (20). In this work, we further investigated this phenomenon focusing on the impact of pectin, as it represents complex sugar polymers typically found in the plant primary cell wall and particularly abundant in the middle lamella layer (24). We first tested the effect of crude pectin extracted from tobacco root PCWP (referred as cPec, Fig. 1ab for composition and related structure). An 8-fold increase of surfactin production was detected at the early exponential growth phase (OD_600_=0.2-0.25) in *B. velezensis* GA1 liquid cultures supplemented with cPec compared to an un-supplemented culture (Fig. 1cd). Surfactin production was also 10 times enhanced upon addition at the same concentration of pure commercially available homogalacturonan (HG) with high degree of polymerization (DP) (Fig. S1ab) but low level of methyl-esterification (HGLM) according to the manufacturer (Fig. 1d). HG was tested as the most abundant pectic polysaccharide constituent, which represents 65% of crude primary cell wall pectin (24). Production of this lipopeptide was also enhanced to a similar level upon addition of highly methylated HG (HGHM), showing that the degree of methyl-esterification of the polymer is not a major trait influencing perception by the bacterium (Fig. S2). Altogether, this supports a key role of the pectin backbone as plant molecular pattern that is sensed by the bacterium to stimulate surfactin synthesis.

**Figure 1:**
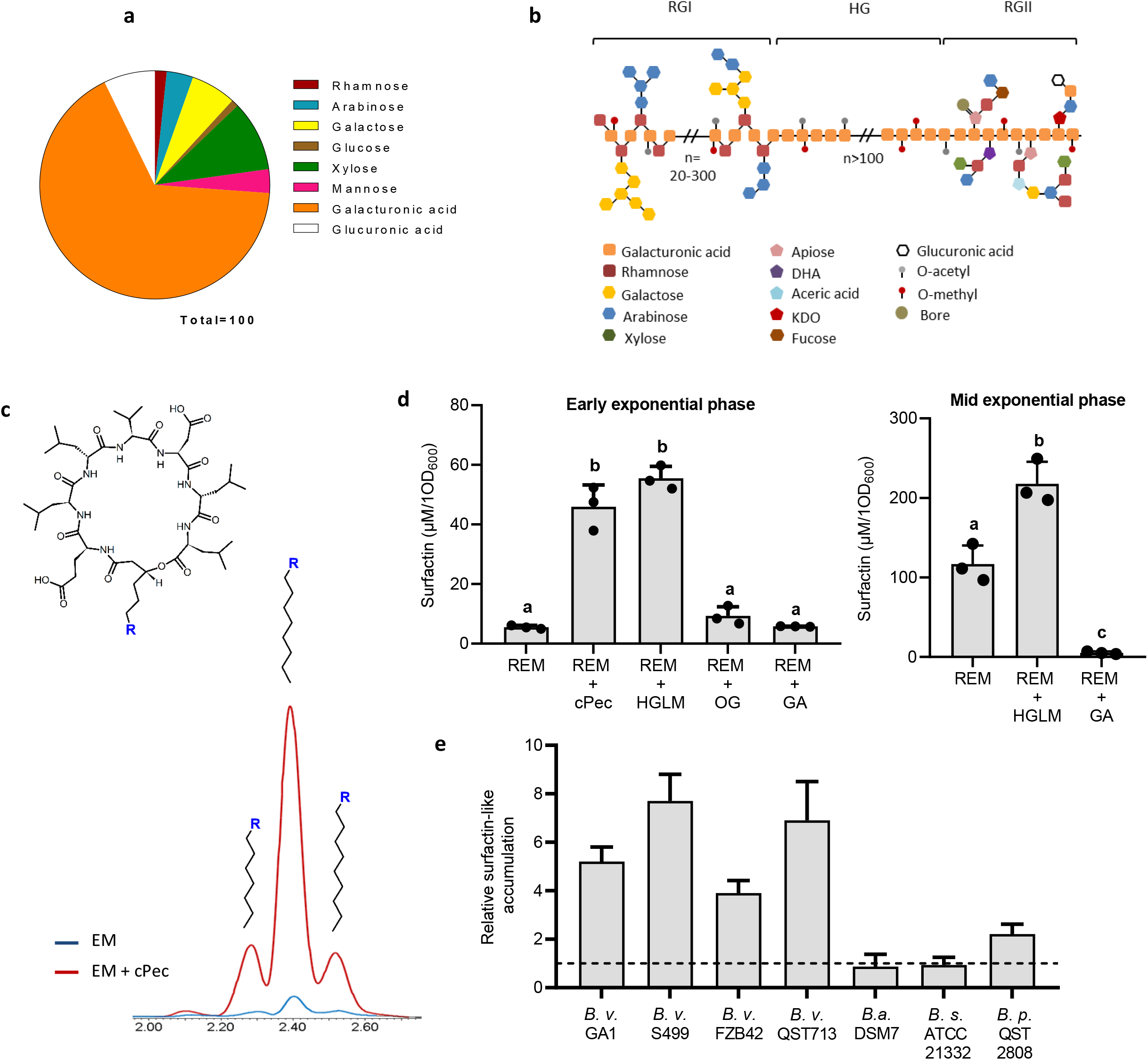
Impact of pectin on early surfactin production. **a** Sugar composition analysis of crude pectin (cPec) extracted from tobacco roots. Composition is expressed as Molar Ratio percentage (Molar %) for each fraction. Galacturonic acid (orange) constituting the pectin backbone (**b** for schematization) is the main sugar of the cPec fraction. Other minor sugars (rhamnose, galactose, arabinose…) are typically found in the pectin side chains (Mohnen et al. 2008, **b**). **b** Schematization of pectin structure. Homogalacturonan (HG) contains an assembly of at least 100 galacturonic acid (GalA) residues that can be acetyl or methyl esterified. Rhamnogalacturonan I (RGI) is constituted by a succession of GalA-Rha dimers, each one containing an alternance of rhamnosyl and galacturonic acid units. The Rha unit can be branched with variable neutral sugar side chains including essentially galactosyl and/or arabinosyl units. Rhamnogalacturonan II (RGII) structure is well conserved within the HG polymer. RGII englobes 9 GalA units substituted by four side chains with complex sugars, including apiose, DHA, aceric acid and KDO, neutral sugars like, rhamnose, galactose, arabinose, xylose, and fucose or also organic acids such as galacturonic and glucuronic acid. RGII can also complex with Bore allowing a crosslink between two HG molecules. **c** Surfactin (cyclic structure represented up) production in a root exudates mimicking (REM) medium at early growth phase (OD_600_=0.2) with (red chromatogram) or without (blue chromatogram) addition of crude pectin extract added to the GA1 cultures. The main peak represents C15 surfactin whereas the minor left and right peaks represents C14 and C16 surfactins, respectively. **d** Surfactin accumulation in the early (left panel, OD_600_=0.2) and mid (right panel, OD_600_=0.35) exponential growth phase of GA1 cultures in REM medium supplemented with different sized pectin fragments : homogalacturonan low methylated (HGLM), DP>150; oligogalacturonides (OG), DP=15; galacturonic acid (GA), DP=1. Means ± std err. from three biological replicates of one experiment are shown. Significate difference between each condition is indicated by different letters, p-value < 0.01. **e** Comparison of surfactin induction level by HGLM in the early exponential growth phase for different *Bacillus* species : *Bacillus velezensis* (*B; v*), *Bacillus amyloliquefaciens* (*B. a*), *Bacillus subtilis* (*B. s*) and *Bacillus pumilus* (*B. p*). For each strain tested, surfactin accumulation was normalized with the control condition without HGLM represented by the black dotted line. Means ± std err. from three biological replicates are shown.

Interestingly, by screening the CAZy database (25) for genes encoding carbohydrate-active enzymes potentially involved in PCWP degradation by *B. velezensis*, two putative pectate/pectin lyases encoding genes were detected. These two genes, referred as *pelA* and *pelB* (accessions *GL331_08735* and *GL331_04125 in B. velezensis* GA1, respectively), are highly conserved among all sequenced *Bacillus* genomes that belong to the “Operational Group *B. amyloliquefaciens*” (Table S1) (26). *pelA* and *pelB* are readily expressed in GA1 *in vitro* and the corresponding enzymes efficiently convert HG into unsaturated oligogalacturonides with consistent activity occurring at the beginning of stationary phase (Fig. S2). However, the bacterial perception of oligomers with lower polymerization degree compared to HG is not obvious since oligogalacturonides (OG) did not stimulate surfactin biosynthesis (Fig. 1d, Fig. S1c for OG characterization). Supplementation with galacturonic acid (GA) led to a reduction of surfactin production at mid exponential phase (OD_600_=0.35, Fig. 1d). Surfactin production is thus specifically boosted upon sensing long degree of polymerization (DP) polymers, but is somehow inhibited in presence of GA constituting the pectin backbone. Such HG-driven surfactin stimulation also occurs in other *B. velezensis* isolates tested (FZB42, QST713 and S499) and to a lower extent *B. pumilus* QST 2808. It does not occur in the non-rhizosphere dwelling isolates *B. amyloliquefaciens* DSM7 or *B. subtilis* ATCC 21332 (Fig. 1e) suggesting that this trait may be specific to bacilli with a plant-associated lifestyle.

### The root nutritional context favors early surfactin production

*Bacillus velezensis* quickly colonize tomato plantlets in a gnotobiotic system and forms visible biofilm-like structures covering the main root and embedding lateral roots after 24-48h post inoculation (Fig. 2a). This is correlated with consistent *srfAA* gene expression and surfactin production rate in the cell population at these early times but it was maintained albeit to a lower level, over the investigated timeframe of seven days (Fig. 2ab). Since surfactin enhancement linked to the perception of the pectin backbone is only transient (Fig. 1d), we hypothesized that root exudates, constantly secreted by the plant, may also positively impact the synthesis of the lipopeptide. Surfactin production rate was thus compared upon growth in a classical laboratory medium (LB) and in a root exudate-mimicking medium (REM) reflecting the content of carbohydrates typically released by tomato or tobacco roots (27). It revealed an earlier and higher production by cells growing in REM (Fig. 2c). Surfactin production in REM medium is initiated earlier and is more efficient in *B. velezensis* compared to other closely related but non plant-associated species such as *B. amyloliquefaciens* or *B. subtilis* (Fig. 2d).

**Figure 2:**
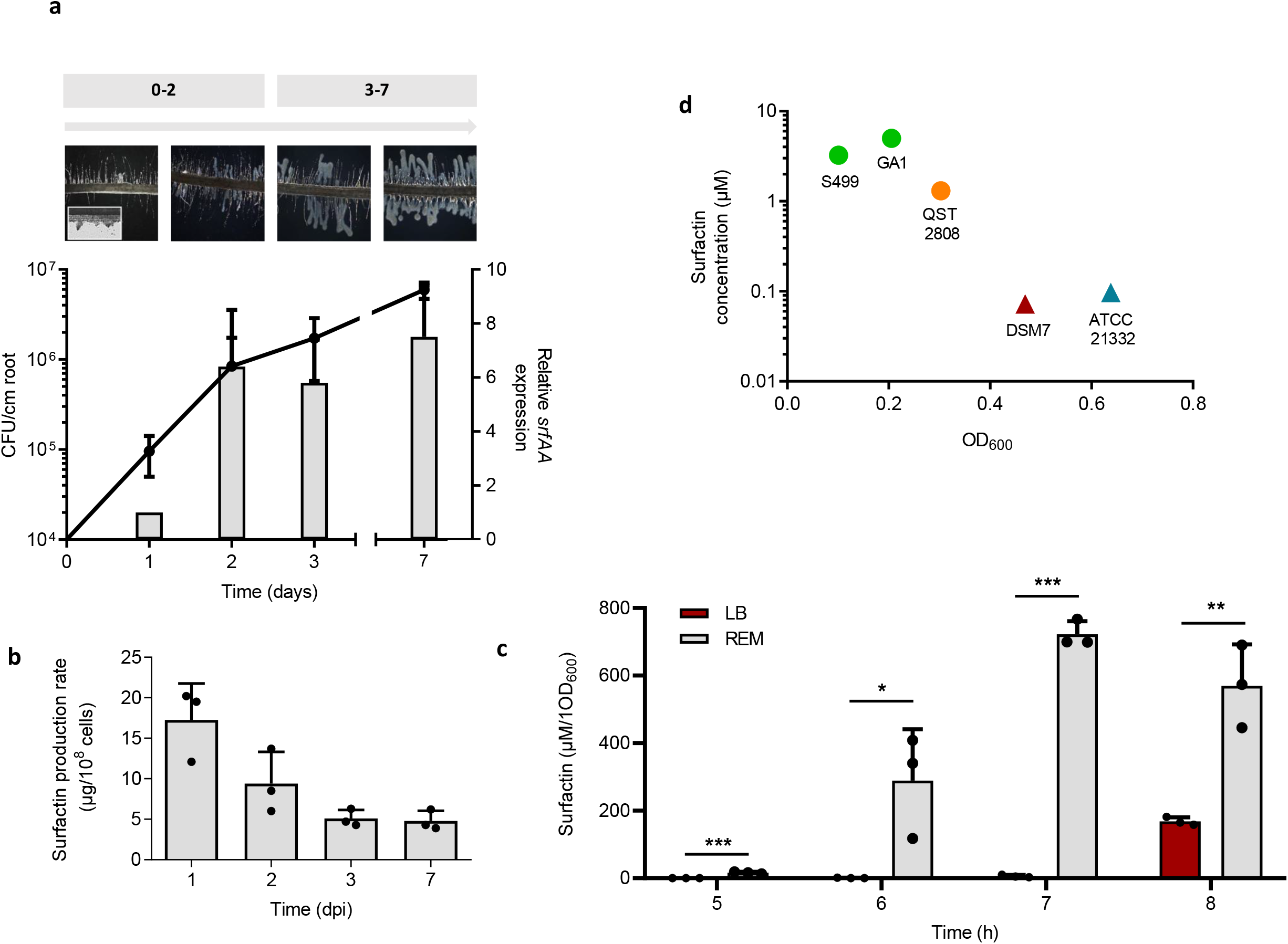
Impact of the specific rhizosphere nutritional context on early surfactin production. **a** Evaluation of bacterial population (black line, left axis) and relative *srfAA* expression on roots (grey bars, right axis) in a time frame of seven days post inoculation (dpi). *Bacillus* progression on roots characterized by a biofilm formation was assessed by microscopy at each time point (upper part). **b** Surfactin production rate on roots. Means ± std err. from three biological replicates of one experiment are shown **c** Surfactin accumulation measured by UPLC-MS in a 8h time course experiment in REM medium (grey bars) compared to LB medium (red bars). Means ± std err. from three biological replicates of one experiment are shown *** P-value <0.001, ** P-value <0.01, * P-value <0.05 **d** Comparison of early surfactin accumulation (µM of surfactin on y axis linked to OD_600_ on x axis) in different *Bacillus* species, including *B. velezensis* (GA1 and S499 in green), *B. pumilus* (QST 2808 in orange), *B. amyloliquefaciens* (DSM 7 in red) and *B. subtilis* (ATCC 21332 in blue). Circle symbols are representing plant associated bacteria whereas triangle symbols are representing non-plant associated bacteria.

Addition of HG in REM medium compared to LB revealed a cumulative effect of this PCWP and root exudates on surfactin production (Fig 3a). This could be of clear ecological benefit for the bacterium since surfactin is known to favor motility of multicellular communities and biofilm formation (16, 28, 29). However, a recent study questioned the real role of surfactin in these key functions, since it production appears as non-essential for pellicle biofilm formation in *B. subtilis* NCIB 3610, suggesting a strain dependant role (30). We previously reported that motility and biofilm formation are boosted upon growth on root exudates (27). Here we show that HG supplementation also favors *B. velezensis* GA1 spreading on low-agar medium (Fig 3b) and early biofilm formation based on pellicle development at the air-liquid interface (31) (Fig 3c). The role of surfactin in swarming, pellicle formation and early root colonization was further confirmed for *B. velezensis* GA1. Indeed, swarming motility on low agar plates was almost reduced to zero in a surfactin deficient mutant, and the same mutant was more than 3 times less efficient to produce pellicles at the air liquid interface and to promptly colonize tomato roots after 1 day post inoculation when compared to the WT (Fig 3def). Collectively, these data allow correlating the positive impact of PCWP on bacterial motility, biofilm formation and early root colonization through an anticipated surfactin production in *B. velezensis*.

**Figure 3:**
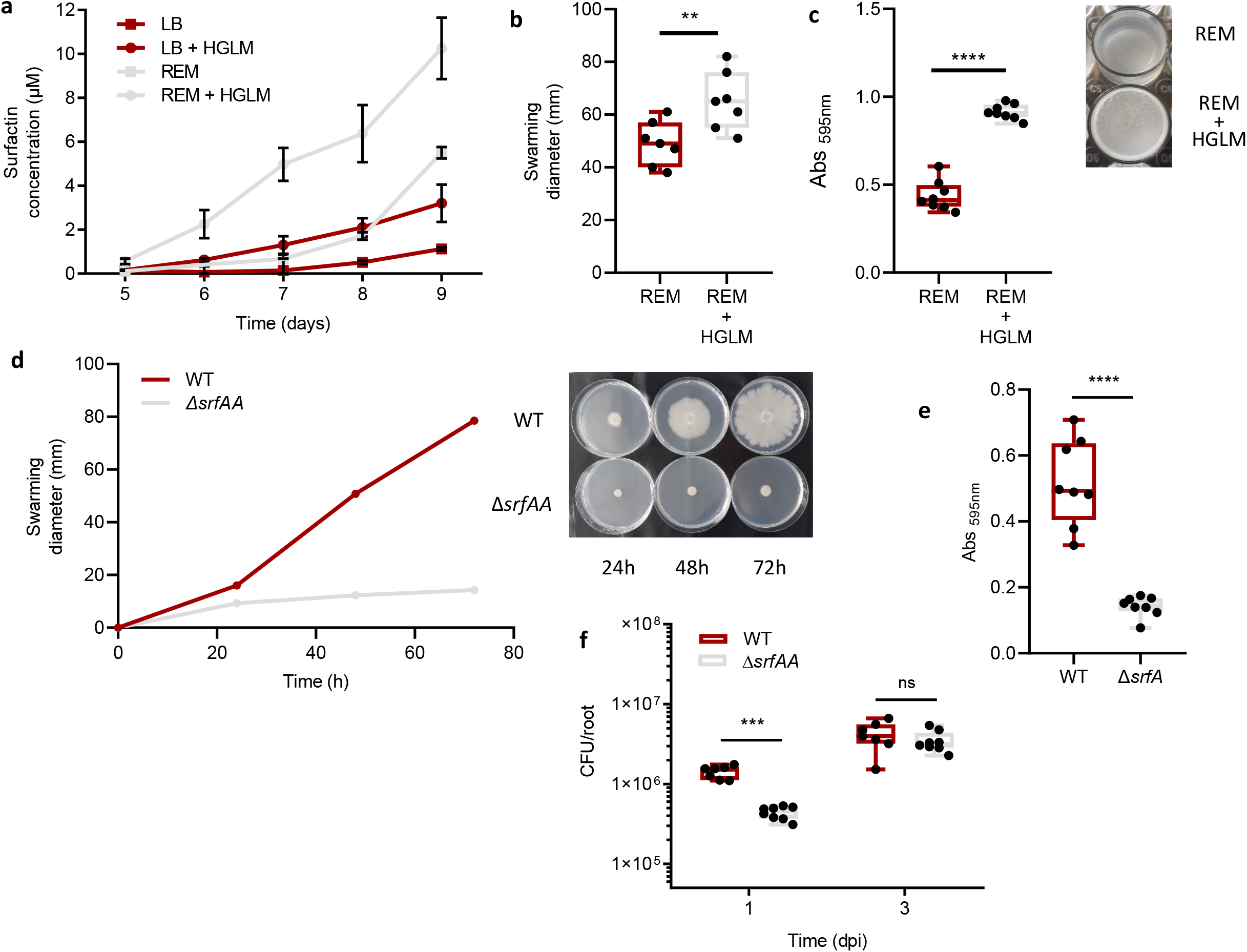
Ecological importance of an early surfactin accumulation. **a** Evaluation of HGLM and root exudates synergistic effect on early surfactin production. Time course experiment for surfactin quantification was performed in REM (grey curves) and LB (red curves) medium with (circle symbols) or without (square symbols) addition of HGLM. Means ± std err. from three biological replicates of one experiment are shown **b** Swarming potential of *B. velezensis* GA1 on soft agar plates after addition of HGLM or not. The box plots encompass the 1^st^ and 3^rd^ quartile, the whiskers extend to the minimum and maximum points, and the midline indicates the median (n=7 biological replicates of one experiment). **c** Evaluation of *B. velezensis* ability to form pellicles on microwells plates after addition of HGLM or not. The box plots encompass the 1^st^ and 3^rd^ quartile, the whiskers extend to the minimum and maximum points, and the midline indicates the median (n=8 biological replicates of one experiment). Pellicle formation is illustrated on the right **d** Comparison of *B. velezensis* GA1 WT and (red) and a Δ*srfAA* mutant (grey) for their swarming potential in a time course study. Means ± std err. from three biological replicates of one experiment are shown. Time course study is illustrated right. **e** Comparison of pellicle formation between GA1 WT strain (red) and a Δ*srfAA* mutant (grey). The box plots encompass the 1^st^ and 3^rd^ quartile, the whiskers extend to the minimum and maximum points, and the midline indicates the median (n=8 biological replicates of one experiment) **** P-value <0.0001. **f** *In vitro* comparison of root colonization ability of GA1 (red boxes) and GA1 Δ*srfAA* (grey boxes) on tomato plantlets. The box plots encompass the 1^st^ and 3^rd^ quartile, the whiskers extend to the minimum and maximum points, and the midline indicates the median (n=7 biological replicates of one experiment) *** P-value <0.001, ns non significant.

### Surfactin induction by PCWP is not linked to major transcriptional changes

Both HG and root exudates stimulate surfactin production in GA1. However, while no activation of the *srfA* biosynthetic gene cluster was observed upon HG addition (Fig. 4a), an early and high surfactin gene expression was measured in *srfAp*::*gfp* cells growing in REM compared to LB medium (Fig. 4b). To unravel transcriptome wide changes in GA1 associated with the perception of HG, RNA-sequencing was performed on cells grown in REM with our without addition of HG and collected at various time points (lag, early exponential and a mid-exponential phases). The data confirmed that HG perception is not linked to an increased expression of the *srfA* operon but also revealed a quite limited and transient transcriptional reprogramming with only 58 genes differentially expressed over this timeframe (Table 1). Remarkably, more than 30% of these genes are involved in stress response or cell wall modifications and are down regulated in the presence of HG (Fig. 4c). We thus hypothesize that a long-term co-evolution process may have facilitated *Bacillus* establishment on the roots by the inhibition of a costly stress response after perception of HG. Addition of HG also leads to a 4.2-fold reduced expression of *flgM* encoding an inhibitor of SigD, the σ factor involved in the activation of motility related genes (32). This may contribute to enhanced spreading of multicellular communities in addition to the positive effect of surfactin mentioned above.

**Table 1:**
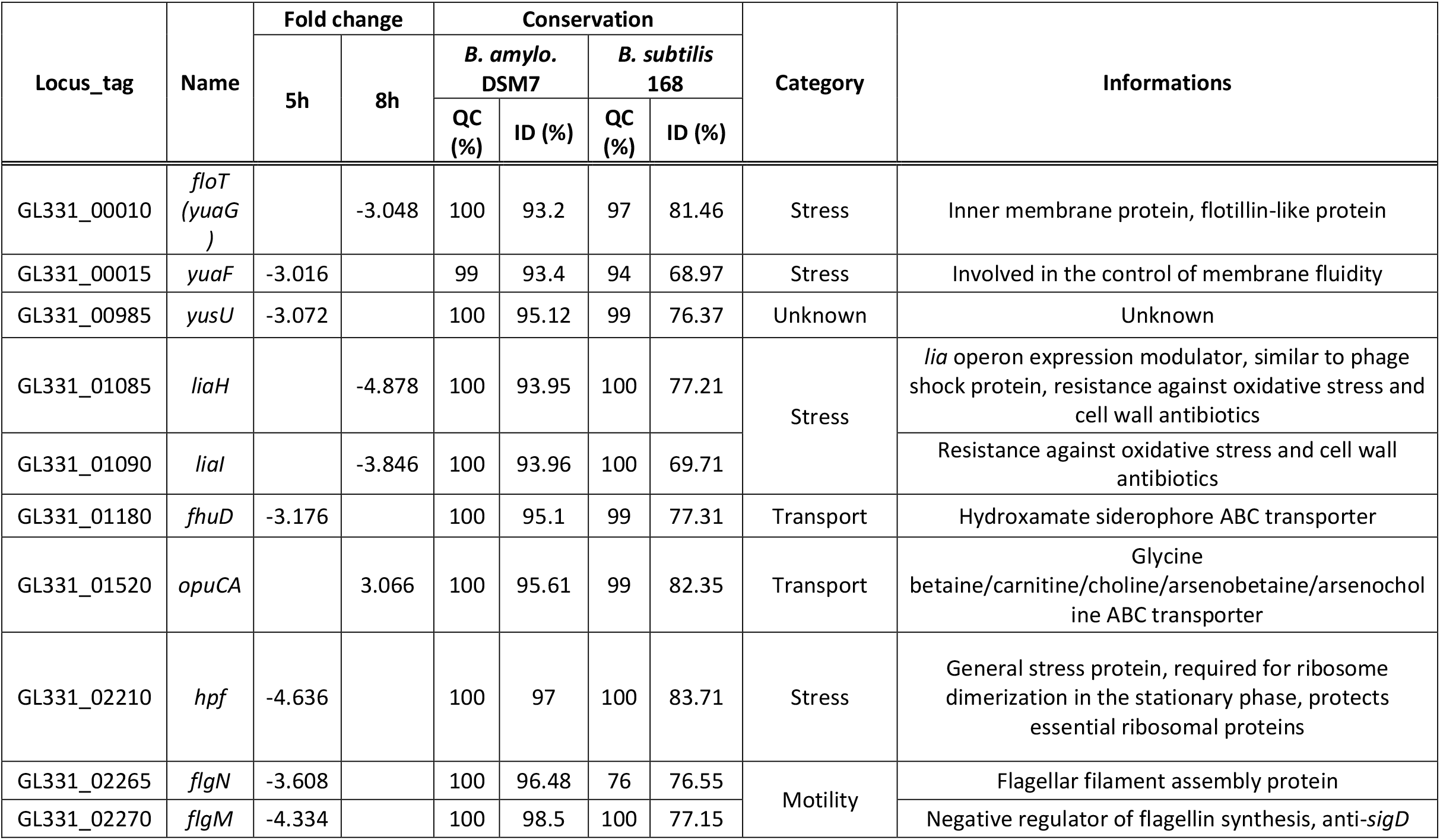
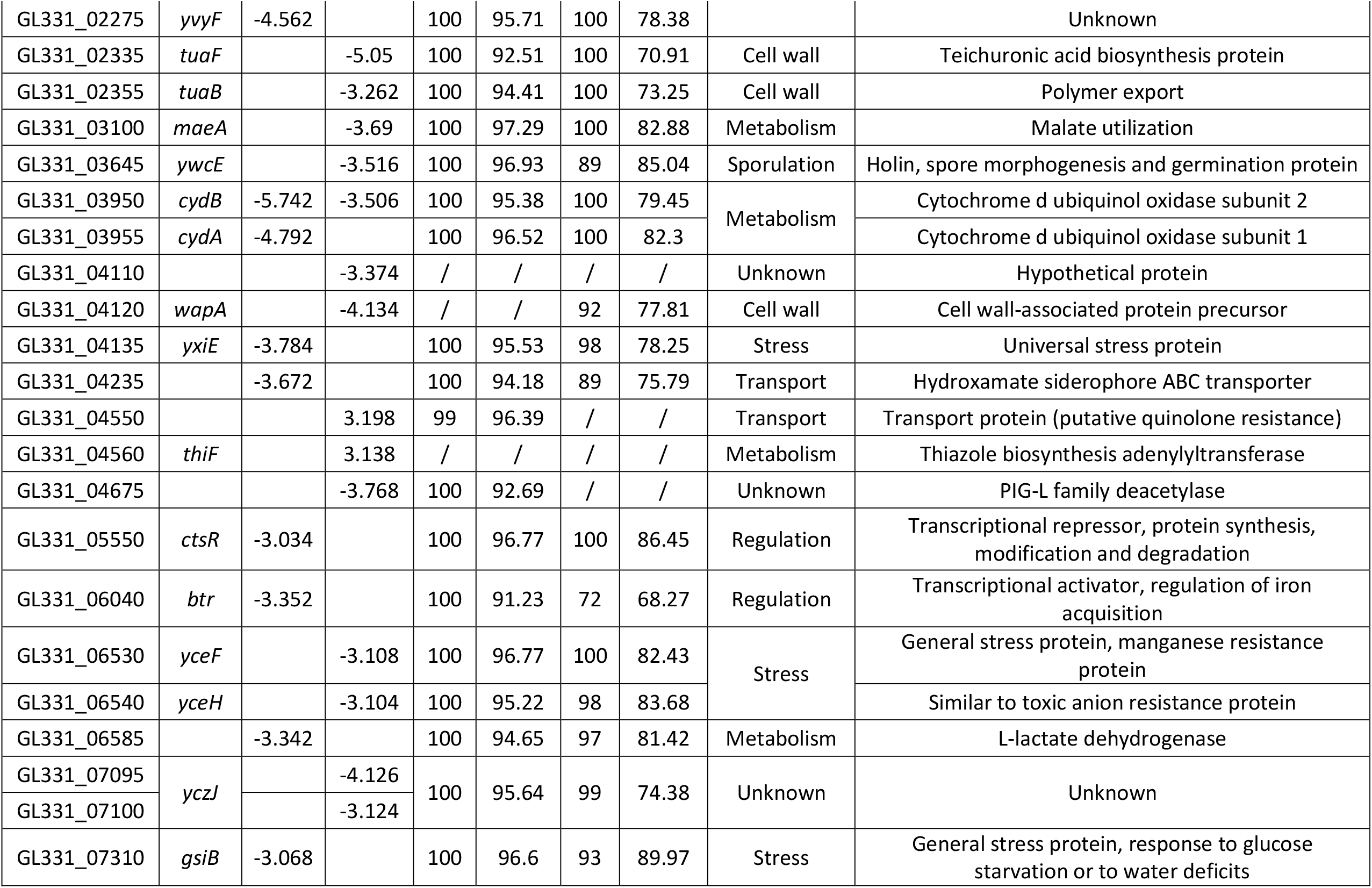
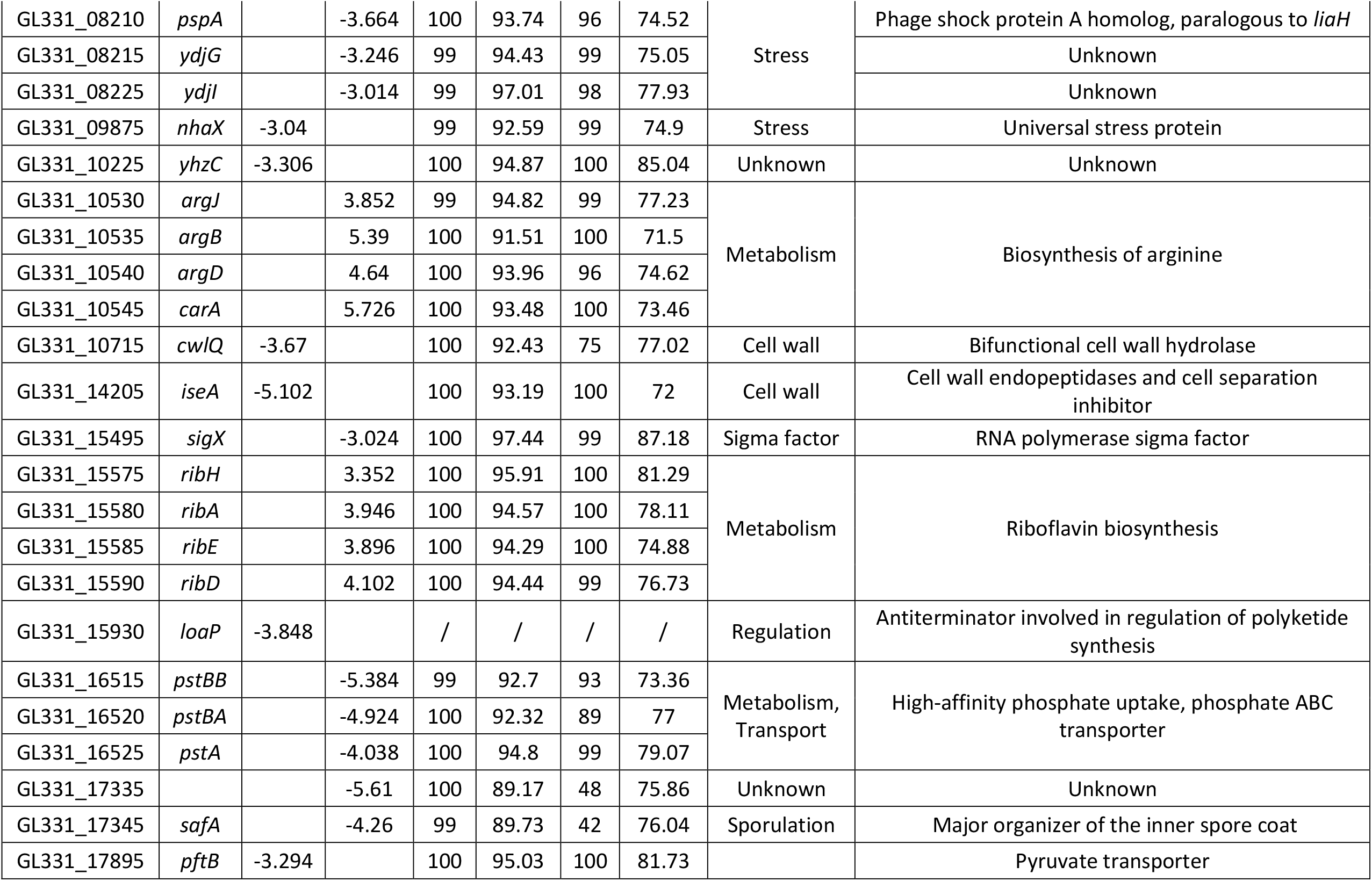
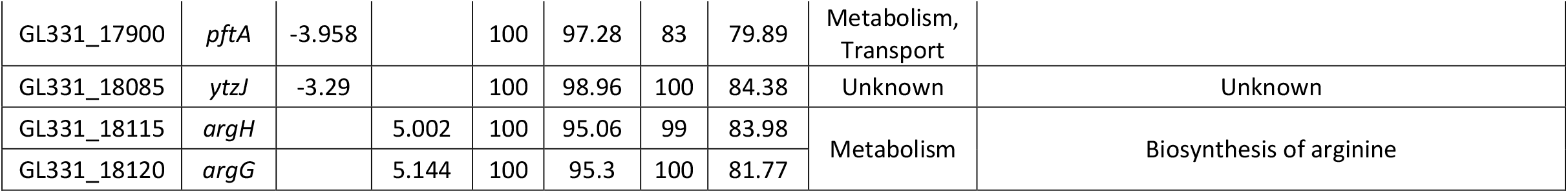
Differentially expressed genes in *B. velezensis* GA1 after HGLM perception

**Figure 4:**
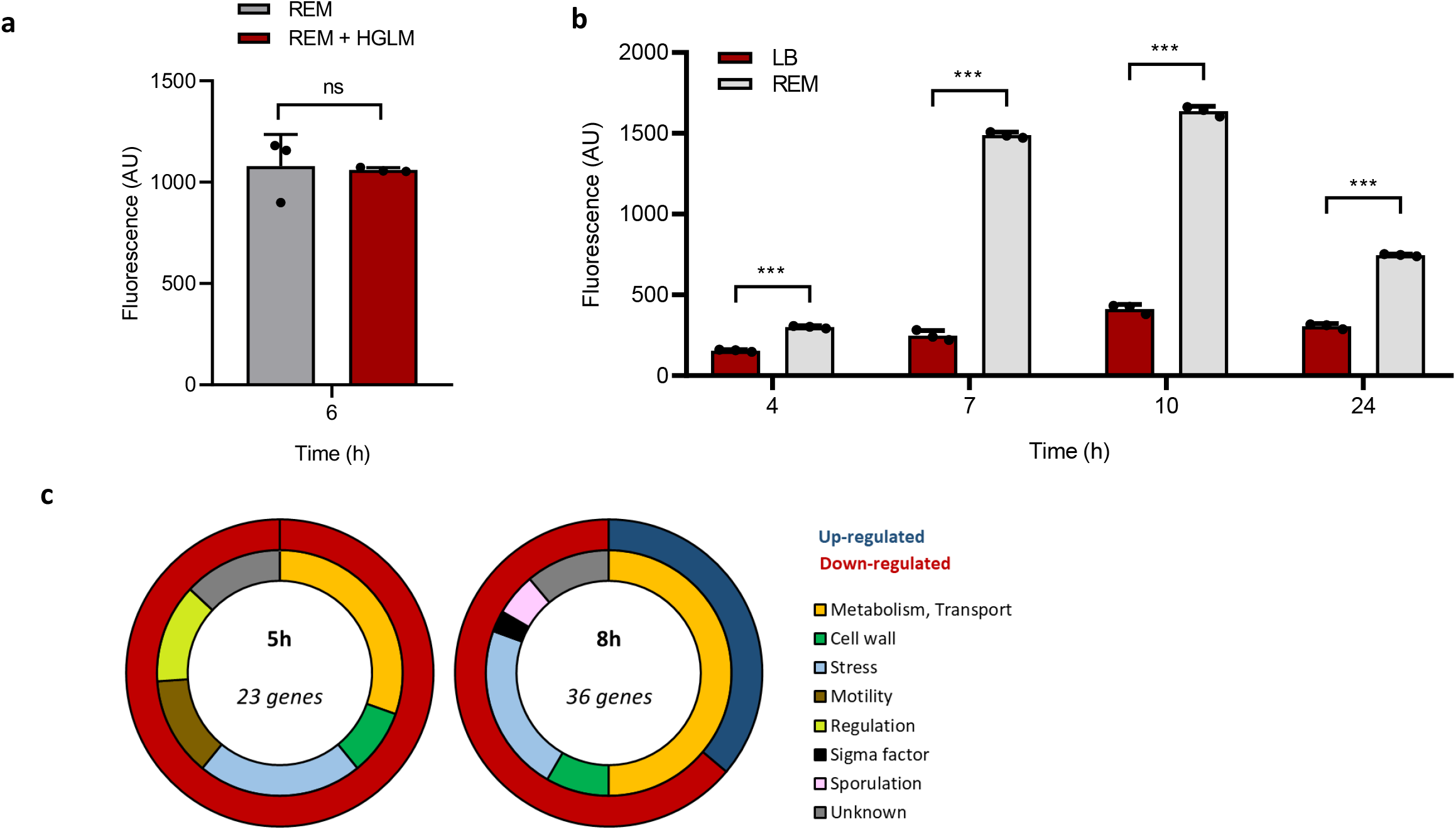
Impact of plant triggers perception on *Bacillus* transcriptome. **a** Surfactin expression measured by fluorescence in the GA1 *PsrfAp::gfp* reporter strain at early exponential phase in REM medium (grey bars) compared to REM medium supplemented with HGLM (red bars). Means ± std err. from three biological replicates of one representative experiment are shown ns = non significate **b** Surfactin expression measured by fluorescence in the GA1 P*srfAp::gfp* reporter strain in a 24h time course study in EM medium (grey bars) compared to LB medium (red bars). Means ± std err. from three biological replicates of one representative experiment are shown *** P-value <0.001. **c** Classification of the different genes carrying a significant fold change (1.5 log2) 5 and 8 hours after addition of HG when compared to the control condition. The outer circle represents the proportion of up (dark blue) and down (red) regulated genes. The inner circle represents the proportion of genes belonging to the different functional family described in the legend.

### Root exudates drive the bacterium to form surfactin homologues with long fatty acid chain (LFAC) and variants enriched in valine

The NRPS machinery works as an assembly line in which each module is responsible for recruiting and binding a specific amino acid to the nascent peptide after a first lipo-initiation step for binding the fatty acid (FA) taken up from the cellular pool (Fig. 1A) (33, 34). In that way, surfactin is typically composed of saturated C_12_ to C_19-_FA of the linear, iso or anteiso type of branching (35). Beside an increased production of surfactin, we also observed an effect on the pattern of surfactin variants synthesized by *B. velezensis* in the presence of artificial plant exudates, as well as in naturally produced exudates and in planta upon root colonization (Fig. S4). Indeed, UPLC-MS profiling revealed that the surfactin pattern produced by GA1 in REM medium is enriched in surfactin *iso-*C_14_ (*i*C_14_) and other variants compared to LB medium (Fig. 5b). They correspond to variants of the canonical structure with substitution of Leu by Val for the last residue of the cyclic peptide moiety (Val_7_) and, to a much lower extent, to the same substitution in position 2 (Val_2_, Fig. 5c, see Fig. S5). Valine is used both as precursor for the synthesis of branched fatty acids with an even number of carbons, and as a building block by the NRPS to form the peptide moiety. Supplementation of the medium with deuterated L-Val-d^8^ resulted in an additional increase in the proportions of surfactin *iso*-C14 and Val_7_ isoforms labeled at the expected positions in the peptide and in the fatty acid tail (Fig. S6). Based on these data, the higher relative proportions of *i*C_14_Val^7^ formed in REM, but also *in planta* (Fig. 5c), most probably result from some enrichment of the intracellular pool in valine upon growth in the presence of root exudates (Supplementary Discussion). Given the reduced specificity of NRPS domains involved in selection and activation of leucine at positions 2 and 7, the megaenzyme would preferably bind valine as it is more available in the pool.

**Figure 5:**
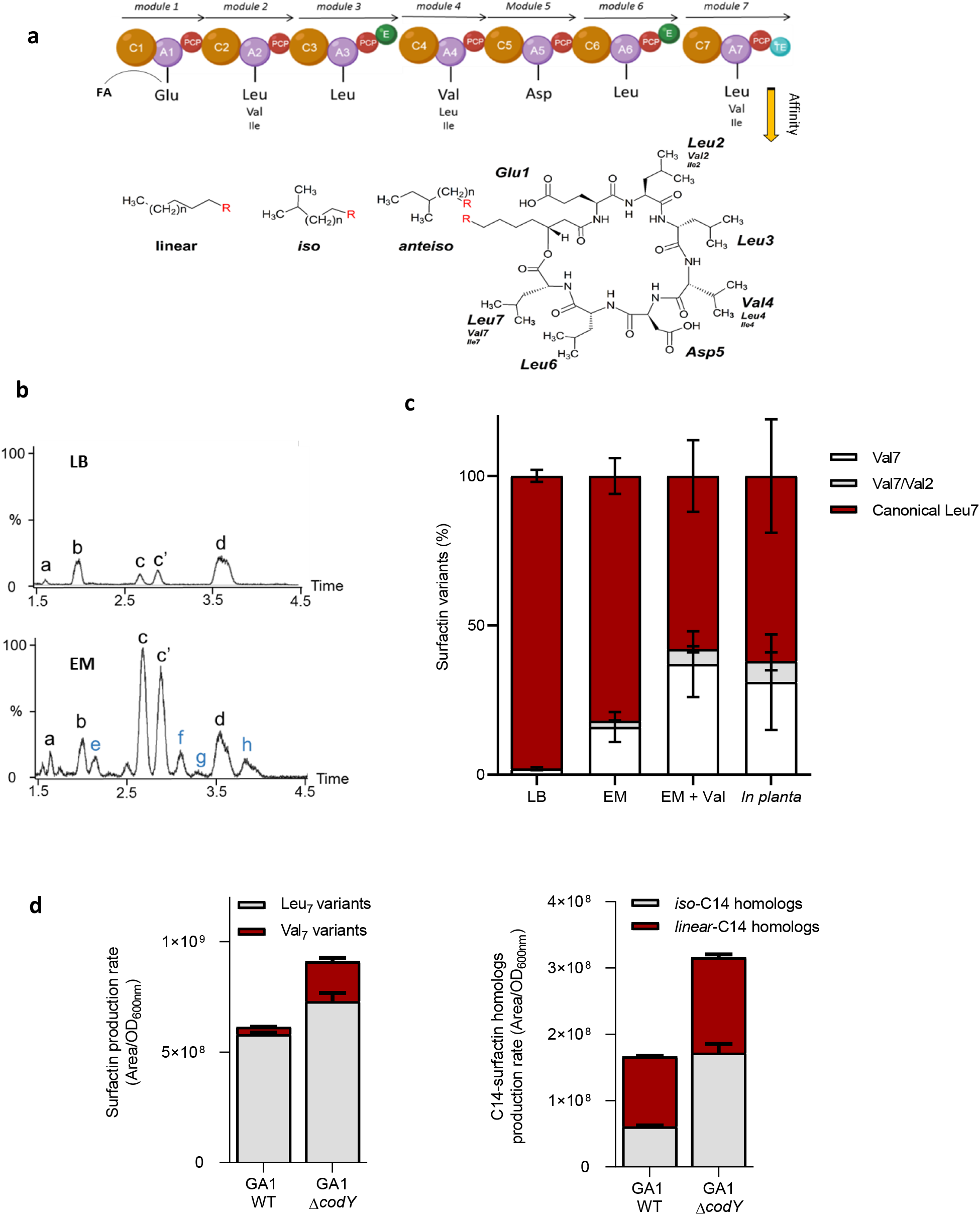
Qualitative impact of root exudates on surfactin production. **a** Representation of the NRPS machinery leading to the assembly of the surfactin molecule. This mega-enzyme is organized in 7 functional units called modules which are each responsible for the incorporation of one amino acid building block into the growing peptide chain. Each module is subdivided into different domains including an adenylation (A, violet circle) and a peptidyl carrier protein (PCP, red circle) catalyzing the peptide initiation, and one condensation domain (C, brown circle) responsible for peptide elongation. The termination of the peptide synthesis is performed by a thioesterase domain (TE, blue circle) in the last module. Modules 3 and 6 also possess an epimerization domain (E, green circle). Surfactin molecule contains a 7 amino acids chain structured as follow: L-Glu – L-Leu – D-Leu – L-Val – L-Asp – D-Leu – L-Leu. In some specific variants, Leu in position 2 and/or 7can be substituted by a Val and more rarely by an Ile, and inversely, Val in position 4 can be substituted by a Leu and also more rarely by a Ile. In addition to the amino acid chain variability, multiple homologs with the same peptidic core but differences in terms of fatty acid chain length (C_12_ to C_17_) or isomerisation of this latter (iso, anteiso or linear configuration) can also be produced. **b** Comparison of surfactin pattern in REM and LB medium. Based on MS-MS analyses, nine different surfactin forms were identified (a: C_12_-Glu-Leu-Leu-Val-Asp-Leu-Leu ; b: C_13_-Glu-Leu-Leu-Val-Asp-Leu-Leu ; c : *iso-*C_14_-Glu-Leu-Leu-Val-Asp-Leu-Leu ; c’: *n-*C_14_-Glu-Leu-Leu-Val-Asp-Leu-Leu ; d : C_15_-Glu-Leu-Leu-Val-Asp-Leu-Leu ; e: C_13_-Glu-Leu-Leu-Val-Asp-Leu-Val ; f: C_14_-Glu-Leu-Leu-Val-Asp-Leu-Val ; g: C_14_-Glu-Leu-Leu-Val-Asp-Leu-Val and h: C_14_-Glu-Val-Leu-Val-Asp-Leu-Val.) **c** Relative proportions of surfactin variants in LB, REM, REM supplemented with valine, and *in planta*. **d** Qualitative and quantitative role of CodY on surfactin production. In a WT strain, 95% of the surfactin molecules are carrying a Leu in position 7 (grey bars) and only 5% are carrying a Val (red bars) whereas in Δ*codY* mutant almost 25% of the surfactin molecules are carrying a Val in position 7 and 75% a Leu. In addition, amount of total surfactin production rate of 150 % can be observed in Δ*codY* mutant compared to WT strain. Proportion of iso-C14 is also affected by CodY, 36 % of total C_14_ are iso-fatty acid (grey bars) and 64% are linear (red bars) in WT strain whereas in Δ*codY* mutant 55% of C_14_ are *iso-*C_14_ and 45 % are linear. Again, total amount of C_14_ is higher in Δ*codY* mutant (increase of 190 %).

As already described in *B. subtilis* (36, 37), the pleiotropic regulator CodY acts as repressor of surfactin synthesis in *B. velezensis* GA1 as illustrated by the 1.9-fold increase in production by the Δ*codY* mutant of strain GA1. Interestingly, CodY activity/*codY* expression is also itself negatively impacted by high cellular concentrations in branched chain amino acids (38). Both quantitative and qualitative changes in surfactin production upon growth in exudates could therefore be, at least partly, due to a lower CodY activity (Supplementary Discussion). In support to the role played by this regulator, a similar impact on surfactin pattern were observed by deleting *codY* in GA1 or by supplementing the culture medium of the wild-type with valine (Fig. 5d).

### Long fatty acid chain surfactins act as key triggers of receptor-independent plant immunity

Based on the potential of surfactin as host immunity elicitor (9, 39), we next wanted to evaluate the possible relevance of quantitative and qualitative modulation of the surfactin pattern driven by the plant for its own benefit.

Upon application as root treatment, pure surfactin used as mixture of isoforms formed in REM medium, induced systemic resistance in hydroponically-grown tobacco plants providing approximatively 45-50% significant disease reduction on leaves subsequently infected with the pathogen *Botrytis cinerea* (Fig. 6a). The various isoforms were then HPLC-purified and tested individually revealing that only long fatty acid homologues (C_14_/C_15_) provided systemic protection to a similar level whereas short fatty acid homologues (C_12_/C_13_) were inactive (Fig. 6b). Moreover, plant immunization by surfactin is dose-dependent and concentrations up to 5 µM are sufficient to significantly stimulate ISR (Fig. 6c). Interestingly, such low µM concentration are actually in the range of those that could accumulate in the root vicinity within a few days upon colonization by GA1 (Fig. S7).

**Figure 6:**
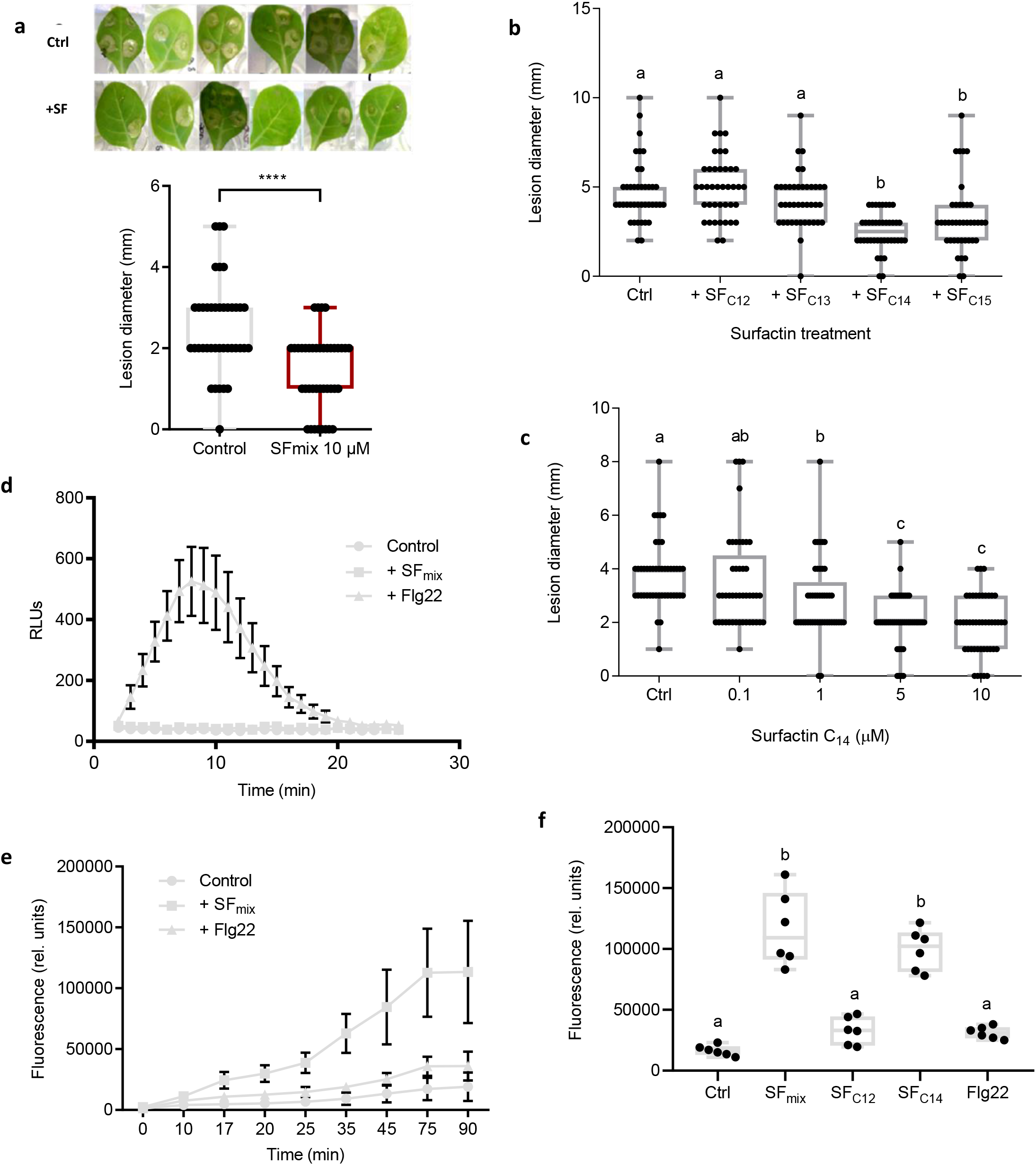
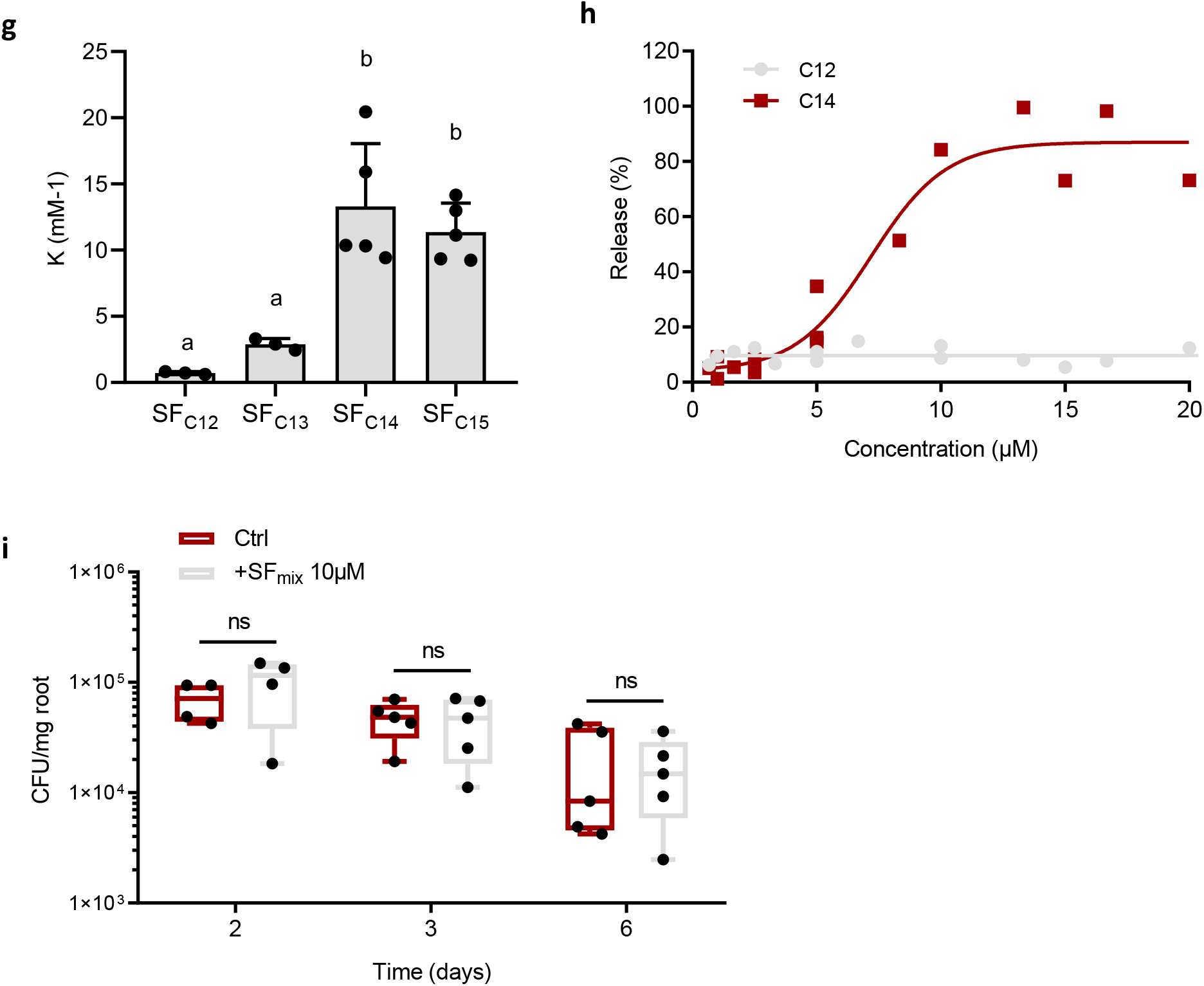
Impact of surfactin homologues on Solanaceae plant immunity. **abc** Systemic resistance induced in hydroponically-grown tobacco by surfactin and expressed as reduction of *B. cinerea* infection (illustration of the reduction in the diameter of spreading lesions on infected leaves) in plants treated at the root level prior to pathogen inoculation on leaves compared to control plants. Data represent results grouped from two independent experiments with similar results and each involving 5 plants with 4 lesions on the second leave (n=40). The box plots encompass the 1^st^ and 3^rd^ quartile, the whiskers extend to the minimum and maximum points, and the midline indicates the median (n=7 biological replicates of one experiment). **a** Effect of surfactin homologues (SF mix) as naturally co-produced by the bacterium (C12/C13/C14/C15 in relative proportions 8/17/33/42%) **** P-value <0.0001 **b** Effect of HPLC-purified surfactin homologues applied at 10 µM with fatty acid chains from C12 to C15. Significate difference between each condition is indicated by different letters, p-value < 0.05 **c** Effect of the most active C14 homologue tested at various concentrations. Significate difference between each condition is indicated by different letters, p-value < 0.05 **de** Stimulation of oxidative burst in root tissues upon treatment with a SF mix and to the response observed by treating roots with flagellin (flg22, 1 µM) used as positive control. **d** Stimulation of apoplastic ROS accumulation (DCFH-DA fluorescent probe) in root tissues upon treatment with a surfactin mix applied at 15 µM. Means and standard deviations are shown for one representative experiment performed on nine samples per treatment each containing three root segments (approx 100 mg FW) collected from different plants (n=9). Similar trend was obtained in an independent assay. **e** Stimulation of cytoplasmic hydrogen peroxide production in root cells. Means and s.d. were calculated from measurements performed on three samples per treatment each containing three root segments (approx 100 mg FW) collected from different plants. Data represent values obtained from two independent experiments (n=6 per treatment). **f** Stimulation of cytoplasmic hydrogen peroxide production in root cells after treatment with C_12_ and C_14_ surfactin homologues as representative of short and long fatty acid chains respectively. Flg22 was used as control. The box plots encompass the 1^st^ and 3^rd^ quartile, the whiskers extend to the minimum and maximum points, and the midline indicates the median (n=6 biological replicates of one experiment).Significate difference between each condition is indicated by different letters, p-value < 0.0001. **g** Binding coefficient (K) of Surfactin homologues (C_12_ to C_15_) to large unilamellar vesicles (LUV) composed by PLPC/Sitosterol/Glucosylceramide (60:20:20 molar ratio). Means ± std err. from three to five biological replicates of one representative experiment are shown Significate difference between each condition is indicated by different letters, p-value < 0.05 **h** Release of 8-hydroxypyrene-1,3,6 trisulfonic acid (HPTS) from PLPC/Sitosterol/Glucosylceramide (60:20:20 molar ratio) LUV, upon addition of surfactin C_12_ or C_14_ at different concentrations. The ordinate shows the amount of HPTS released after 15 min in the presence of the C_12_ or C_14_ as a percentage of the total amount released by Triton X-100. **i** Influence of roots pretreatment with 10µM of surfactin (blue boxes) compared to non-treated roots (red boxes) on *B. velezensis* GA1 root colonization. The box plots encompass the 1^st^ and 3^rd^ quartile, the whiskers extend to the minimum and maximum points, and the midline indicates the median (n=5 biological replicates of one experiment). Ns= non significate.

We next wanted to correlate this systemic protection induced by the lipopeptide with its potential to trigger locally early immune-related events such as the extracellular burst in reactive oxygen species (ROS) involved in defense and signaling in pathogen-triggered immunity (PTI) (40, 41). By contrast with flagellin (epitope Flg22), one of the best characterized Microbe-Associated Molecular Patterns (MAMPs) isolated from bacterial pathogens, treatment with surfactin did not induced burst in apoplastic ROS in root tissues (Fig. 6d). However, surfactin-mediated ROS signaling still occurs since a clear cytoplasmic ROS accumulation was observed (Fig 6e). Little information is available about the spatio-temporal dynamics of such ROS burst but it may originate from different organelles and has been occasionally described in response to perception of biotic and abiotic stresses (42, 43). Using cytoplasmic ROS as marker, the same trend as for ISR tests could be observed regarding the influence of the structure on the activity of surfactin since long fatty acid homologues but not short ones efficiently stimulated early immune reaction (Fig 6f). This means that a single additional methylene group in the fatty acid tail of the molecule (C_14_ versus C_13_) likely determines its immunization potential (Fig 6b,f). By contrast, substitution of Leu^7^ by a Val in the C_14_ homologue does not impact activity suggesting that the peptide moiety is not essential for perception by plant cells. In addition, the µM concentrations required for optimal eliciting activity of surfactin are very high compared with PAMPs active in the nM range (44). Our previous data showed that surfactin elicitation is still active after treating cells with proteases or after a first application indicating that there is no saturation of high-affinity but low abundance binding sites on receptors (45, 46). All this indicates that surfactin is perceived by plant cells via a mechanism independent of high-affinity pattern-recognition receptors (PRRs) involved in MAMP perception (40, 41, 44, 47, 48). We therefore postulated that surfactin perception relies on some interaction with the lipid phase of the plant plasma membrane. Binding experiments via isothermal titration calorimetry and leakage assays based on the release of fluorescent probe were performed using liposomes prepared with lipids specific to plant plasma membrane (PLPC/sitosterol/glucosylCeramide). It revealed that long fatty acid homologues have a higher affinity for these vesicles than the short fatty acid forms and display a higher destabilizing effect on the lipid bilayer when added at concentrations of 5 µM or higher (Fig. 6gh). These biophysical data thus correlated well with the contrasting biological activities of longer C_14_/C_15_ and shorter C_12_/C_13_ surfactin homologues.

According to the priming concept (49), we previously showed that ISR triggered by the lipopeptide in that plant as well as in tobacco and *Arabidopsis*, is not associated with a fast and strong expression of defensive mechanisms before pathogen infection (20, 39). In order to verify that surfactin elicitation does not cause a massive release of antimicrobials from plant tissues, tomato roots were pre-treated with the lipopeptide before inoculation with *B. velezensis*. As expected, it did not impacted subsequent colonization in terms of rate and dynamics compared to untreated plants indicating the absence of potential adverse effects on the bacterial partner (Fig. 6i).

## Discussion

A large part of the interactions between bacteria and plants is known to be mediated by small-size secreted products (50). However, a better understanding of the chemical cross-talk at the plant-bacteria interface and its impact on bacterial ecology, plant fitness and immune responses remains challenging. In epiphytic soil bacilli, root exudates induce expression of an array of genes involved in various functions such as chemotaxis and nutrient acquisition (51–53). Our data further illustrate that utilization of this cocktail of molecules released by roots but also the perception of some cell wall polymers may also drive these bacteria to efficiently produce key components of the secondary metabolome and more specifically the multifunctional surfactin lipopeptide (20). As an amphiphilic molecule and powerful biosurfactant, surfactin is presumably viewed as membrane active compound with potent antimicrobial activity. However, this lipopeptide is poorly antibacterial and antifungal (54). In *B. velezensis*, more obvious ecological functions of this CLP are to contribute to motility, biofilm formation and roots colonization. An enhanced production upon host perception thus constitutes a major force driving successful rhizosphere establishment.

Homogalacturonan acts as a cue to enhance surfactin secretion by bacterial cells but no transcriptional induction of the corresponding biosynthesis operon was observed. Surfactin synthesis is integrated in a complex network involving several pleiotropic regulators acting directly or indirectly on the expression of the *srfA* operon (55–58). However, we hypothesize that surfactin induction by HG may rather rely on post-transcriptional changes as reported for the effect of the DegU and YczE regulators on production of another CLP, bacillomycin D (59). Despite the relatively close genetic proximity of the two species, our data suggest that regulation of surfactin could be slightly different in *B. velezensis* and *B. subtilis*. As it represents a key infochemical devoted to cross-talk with the host plant, surfactin regulation may have been fine-tuned in rhizosphere species to better fit with the nutritional or more broadly ecological context.

Deciphering the mechanism by which *B. velezensis* recognizes pectin and enhances surfactin production would help to identify candidate genes and pathways that are responsible for plant sensing, ensuring persistence on roots which globally remains very poorly known for beneficial rhizobacteria. We are currently investigating whether some cell surface proteins may act as receptors for homogalacturonan perception and binding as recently described for *Sphingomonas* sp. (60), another beneficial species living in association with plants (61). Some insights could be obtained by scrutinizing the few genes conserved in *B. velezensis* but missing in non-plant-associated *B. amyloliquefaciens* strains that are not responsive to pectin (62). Interestingly, shorter fragments of HG and galacturonic acid do not stimulate surfactin secretion. It is therefore tempting to hypothesize that sensing unaltered polymer could indicate a healthy host suitable for bacterial colonization while the perception of monomers or low DP oligomers may reflect a dead or infected plant that is unable to adequately provide resources.

Our data illustrate for the first time that *B. velezensis* can also modulate qualitatively its surfactin pattern by growing in its natural nutritional context, *i*.*e*. on root exudates. Substitution of leucine by valine in the peptide part is not expected to impact the contribution of the lipopeptide to colonization by the producing strain itself considering the minor effect of these structural changes on motility and biofilm formation potential (18). Small modifications in the peptide sequence may nevertheless avoid surfactin hijacking for use as signal prompting heterologous biofilm formation by closely related competitor species (18). Based on our observations, the most obvious benefit of an increased proportion of long fatty acid chain homologues is for the host plant since they represent the most active forms for priming immunity with no impact on host fitness (20, 39), by contrast with PTI (63, 64). As the bacterial partner does not have to face strong defensive responses from this reaction, it ensures positive mutualistic co-habitation allowing establishment of populations on roots. Persistence of threshold populations is necessary for consistent production of other specialized secondary metabolites more directly involved in warding off both microbial competitors and plant soilborne pathogens in the context of biocontrol.

Surfactin stimulation upon sensing host molecular patterns may thus reflect an aspect of plant-*Bacillus* coevolution as it makes a shared good out of this multifunctional lipopeptide. To some extent, it might represent a facet of the plant-driven selection process resulting in active recruitment of this bacterium as species that provides beneficial functions. Other bacterial genera such as *Pseudomonas* also prevailing in the rhizosphere microbiome actively produce CLPs with similar roles as surfactin. Evaluating whether their synthesis is also modulated by plant cues would conceptually allow broadening the significance of these lipopeptide-mediated inter-kingdom interactions for bacterial ecology, plant health and biocontrol.

## Materials and Methods

### Bacterial media and growth conditions

Cultures were performed at 26°C in root exudates mimicking medium (EM) (27) or in LB medium. To test the effect of plant cell wall polymers, each specific plant polysaccharide was added at a final concentration of 0.1% in the culture medium. Low (HGLM, <5%) and high (HGHM, >95%) methylated homogalacturonan were provided from Elicityl Oligotech whereas oligogalacturonides and D-galacturonic acid were provided from Sigma.

### Strains construction

All the bacteria strains used in this study are listed in table 2. All the primers used in this study are available upon request. To follow the expression level of the *srf* operon in GA1, we constructed a *gfp* transcriptional fusion under the control of the *srf* promoter and integrated it into the *amyE* locus. First, a GA1 *amyE* amplicon containing a native *KasI* restriction site was integrated in the PGEMT easy. In parallel, a *cat*-*gfp* cassette containing respectively (i) a chloramphenicol resistance gene (cat) and (ii) a promoterless *gfpmut3*.*1* gene was amplified with primers containing *KasI* sites at their 5’ extremities using the pGFP star as a matrix (65). The pGEMT *amyE* plasmid and the *cat*-*gfp* amplicon were both digested by *KasI* (NEB) and the two linear fragments with compatible 5’ overhangs were ligated together to obtain the PGEMT *amyEup-cat*-*gfp-amyEdw* plasmid. To construct the final mutation cassette, an overlap extension PCR was assessed by following the method developed by Bryksin and Matsumura (66). One first fragment containing the upper *amyE* homologous region and the *cat* gene, and one second fragment englobing the *gfpmut3*.*1* gene and the lower *amyE* homologous region were both amplified using the PGEMT *amyEup-cat*-*gfp-amyEdw* plasmid as a matrix. A third fragment was amplified using GA1 genome as matrix with chimeric primers designed to obtain a *srf* promoter amplicon flanked by 20 bp connectors in 5’ and 3’ containing respectively homologies to the upper and lower *amyE* fragments. All three fragments were joined together with a second PCR race to obtain the final cassette. *B. velezensis* GA1 transformation was performed after modification from the protocol developped by Jarmer *et al*. (67). Briefly, one colony was inoculated into LB liquid medium at 37°C (160 rpm) during 6h and cells were washed two times with peptone water. Until 1µg of the recombinant cassette was added to the GA1 cells suspension adjusted to an OD_600nm_ of 0.01 into MMG liquid medium (19 g l-1 K_2_HPO_4_ anhydrous; 6 g l-1 KH_2_PO_4_; 1 g l-1 Na_3_ citrate anhydrous; 0.2 g l-1 MgSO_4_ 7H_2_O; 2 g l-1 Na_2_SO4; 50 µM FeCl_3_ (sterilized by filtration at 0.22 µm); 2µM MnSO_4_; 8 g l-1 Glucose; 2 g l-1 L-glutamic acid; pH 7.0). Cells were incubated at 37°C with shaking, and colonies who integrated the cassette by a double crossing over event were selected on LB plate supplemented with chloramphenicol. Proper integration of the *cat*-*gfp* locus was verified by PCR. Knock-out mutant strains were constructed by gene replacement by homologous recombination. A cassette containing a chloramphenicol resistance gene flanked respectively by 1 kb of the upstream region and 1 kb of the downstream region of the targeted gene was constructed by a three partners overlap PCR. This recombination cassette was also introduced in *B. velezensis* GA1 by inducing natural competence as described above (67). Double homologous recombination event was selected by chloramphenicol resistance. Deletion was confirmed by PCR analysis with the corresponding upstream and downstream primers.

**Table 2:** Strains used in this study

| Strain | Characteristics | Sources |
| --- | --- | --- |
| *Bacillus velezensis* |  |  |
| *B. velezensis* GA1 | Wild-type strain | (83) |
| *B. velezensis* GA1 Psrf_gfp | amyE::Psrf_gfp+chl ; Chl+ | This study |
| *B. velezensis*GA1 ΔsrfAA | ΔsrfAA::chl ; Chl+ | This study |
| *B. velezensis*GA1 ΔcodY | ΔcodY::chl ; Chl+ | This study |
| *B. velezensis* S499 | Wild-type strain | (15) |
| *B. velezensis* FZB42 | Wild-type strain | (13) |
| *B. velezensis* QST713 | Wild-type strain | (84) |
| *Bacillus amyloliquefaciens* |  |  |
| *B. amyloliquefaciens* DSM7 | Wild-type strain | ATCC |
| *Bacillus subtilis* |  |  |
| *B. subtilis* ATCC 21332 | Wild-type strain | ATCC |
| *Bacillus pumilus* |  |  |
| *B. pumilus* QST 2808 | Wild-type strain | (85) |
| *Escherichia coli* |  |  |
| *E. coli* dh5α | Wild-type strain | CGSC |
| *E. coli* dh5α pGEM-T Easy amyE | pGEM-T Easy amyE ; Amp+ | This study |
| *E. coli* dh5α pGEM-T Easy amyEup-cat-gfp-amyEdw | pGEM-T Easy amyEup-cat-gfp-amyEdw ; Amp+ Chl+ | This study |
| *E. coli* dh5α pGFP_Star | pGFP-Star ; Chl+ | This study |

### Fluorescence measurement

Fluorescence accumulation was evaluated thanks to the channel FL1 of a BD accuri C6 flow cytometer (Biosciences) with the following parameters: 20000 events, medium flow rate (35 µl.min^-1^), FSC threshold of 20000.

### Genome sequencing

GA1 genome sequence was reconstructed using a combined approach of two sequencing technologies which generated short paired end reads and long reads. The resulted sequences were then used for hybrid assembly. More precisely, genomic DNA was extracted and purified from *B. velezensis* GA1 using the GeneJET Genomic DNA purification (ThermoFisher scientific). First half of extracted DNA was sent to the GIGA sequencing facility (Liège, Belgium), and use as DNA template for illumina MiSeq sequencing after being prepared using nextera library kit illumina. Sequencing run generated 150 bp paired-end read, which were trimmed and corrected using an in-house python script and SPAdes 3.14 (68) before assembly. The second half of extracted DNA was used to generate long reads with a MinION Oxford nanopore plateform. DNA library was constructed using the Rapid Sequencing kit (SQK-RAD0004, Oxford nanopore). Adapters were trimmed from generated reads with Porechop software (https://github.com/rrwick/Porechop). Trimmed reads were then filtered by size (>500) and Q-score (>10) using NanoFilt implemented in NanoPack (69). Finally, hybrid assembly was performed using hybridSPAdes algorithm implemented in SPAdes 3.14 (70).

### Transcriptome library preparation and sequencing

RNA extraction was performed for each sample using the NucleoSpin RNA kit (Macherey-Nagel). Total RNAs were quantified using Nanodrop (Thermofisher). For sequencing, all samples were sent to the GIGA-genomics platform in Liège. Quality was assessed using the RNA 6000 Nano Chip on a 2100 Bioanalyzer (Agilent). cDNA libraries were prepared employing Universal Prokaryotic RNA-Seq, Prokaryotic AnyDeplete kit (Nugen) according to the manufacturer’s instructions, with . cDNA libraries were quantified and normalized by using the KAPA SYBR Fast Mastermix (Sigma-Aldrich) with P5-P7 Illumina primers according to the manufacturer’s instructions. Prepared libraries were sequenced on a NextSeq 550 device (Illumina) by using the following parameters : paired end, 80 cycles read 1, 8 cycles index, 80 cycles read 2.

### RNA-seq data analysis

The raw RNA-seq reads were trimmed using Trimmomatic v0.39 (71). We performed quality control on the trimmed reads using FastQC v0.11.8 (Babraham Bioinformatics). Trimmed reads were mapped to the GA1 reference genome (see section “genome sequencing” for accession numbers) using BWA-mem v0.7.17 (72) with the following settings: mem -k 50 -B 40 -v 1. At least 95.4% of reads uniquely mapped to the annotated reference genome (Table S2). SAMtools v1.9 (73) was used to generate the BAM files and their indices. To calculate the read counts, the python-based tool HTSeq v0.9 (74) was employed with the following parameters: htseq-count -q -s no -f. The Cufflinks function cuffnorm (75) was used to generate the FPKM (fragments per kilobase of transcript per million mapped reads) tables using the following settings: --compatible-hits-norm --library-norm-method classic-fpkm. Genes with low reads counts (<25) were removed before further analysis. Differential expression analysis was conducted according to the DESeq2 pipeline (10.1186/s13059-014-0550-8) with cut-off parameters of p<0.05 and log2-fold-change > 1.5.

### Motility and biofilm assays

Swarming motility assays were performed according to Molinatto *et al*. 2017 (76). Diameter of the bacterial swarming pattern was measured 48h after inoculation on REM soft agar plates (0.8% agar) supplemented or not with 0.1% HGLM. Quantification of total biofilm was performed by crystal violet staining. Strain of interest was inoculated at a final OD_600_ of 0.1 in a 96 wells microplate containing 200 µl of REM medium supplemented or not with 0.1 % HGLM. The plate was incubated at 30°C during 24h without shaking. Medium and planctonic cells were discarded and wells were washed with PBS. Biofilm pellicle was stained with 0.1% crystal violet during 10 min and washed with PBS. The stained biofilm was dissolved with 30% acetic acid. Absorbance was measured at 595 nm.

### Plant growth conditions and roots colonization assays

For sterilization, tomato seeds were first immersed in a 70% ethanol solution during 2 minutes, transferred in a 20% bleach solution under shaking for 20 minutes and rinsed three times with sterile water. Sterilized tomato seeds were pre-germinated on solid Hoagland medium at 22°C under a 16h/8h night/day cycle. After 4 days, 5µL of cultures containing the strain of interest and calibrated at OD_600_=1 were deposited on the root top. After 1 and 3 days of colonization, roots were harvested, deposited separately in a peptone water solution supplemented with 0.1% of Tween, and vortexed vigorously to tear off the bacterial cells from the roots. Several dilutions were plated on LB media to evaluate the level of colonization. Measurements of surfactin production by GA1 cells colonizing roots were performed on 1×1×0.7 cm pieces of gelified medium containing roots based on the assumption that the produced lipopeptide diffused to a maximal distance of 5 mm from each part of the root and is uniformly distributed over the surface as we previously observed via imaging-MS (77). A 10-fold concentration factor was applied to estimate concentrations around the root surface in order to take into account diffusion constraints in a solid matrix. Surfactin was quantified by UPLC-MS as described below.

### Plant cell wall extraction

Tobacco seeds were sterilized as described above for tomato seeds and deposited on Hoagland plates at 22°C during one week for a successful germination process. Each plantlet was then transferred in a seedholder filled with soft agar and put in Araponics boxes containing the nutritive solution described above. Cell wall extraction was performed on 6 weeks old plants grown at 22°C with a 16h/8h day-night alternance. Roots were harvested, lyophilized and reduced to powder using a Retsch MM400 grinder. 500 mg of powder was resuspended in 40 ml of ethanol 80% at 90°C for 20 min. The insoluble cell wall fraction was recovered by centrifugation and the pellet obtained was washed once with water to obtain the Alcoholic Insoluble Residue (AIR) used for fractionation. AIR was freeze-dried before use in fractionation protocol. Sequential extraction of root cell walls was performed using a protocol derived from Carpita (78) and Silva *et al*. (79). Dry AIR was resuspended in 40 ml water and incubated at 100°C for 20 min. Supernatant was recovered after centrifugation as a soluble pectic fraction (cPEC).

### Monosaccharide composition analysis using HPAEC-PAD

Before monosaccharide composition analysis, cPec fraction was dialyzed during 24h against a large volume of water and freeze-dried. 2 mg of dried fraction material was hydrolyzed in 1 ml of 2M Trifluoroacetic acid (TFA) at 121°C for 90 min. TFA was evaporated under nitrogen gas flux and the hydrolysed dried residue was resuspended in 1 ml water, filtered on 0,2 μm cartridge and stored in vials at 20° before HPAEC-PAD. High Performance Anion Exchange Chromatography with Pulsed Amperometric detection (HPAEC-PAD) was used for neutral and acidic monosaccharide composition analysis using a Dionex DX-500 system (Dionex Corporation) equipped with a Carbopac PA-1 analytical column (4 mm × 250 mm). The elution was performed with a flow rate of 1 mL.min^-1^ in a gradient mode. The gradient for neutral sugars (eluent A: deionized water, eluent B: 160 mM NaOH and eluent C: 200 mM NaOH) was 10% B for 25 min, 100% B for 10 min and finally an equilibration step with 10% B (15 min). The gradient for uronic acid (eluent A: 160 mM NaOH and eluent B: 160 mM NaOH + 600 mM AcONa) was 0% B for 5 minutes, 30 minutes of linear gradient from 0 to 100% B, 100% B for 5 minutes and finally an equilibration step with 0% B (10 minutes). Detection was performed with a pulsed amperometric ED50 detector (Dionex Corporation). 20 mL of sample was injected with an autosampler. Each carbohydrate concentration was determined after integration of the respective areas (Chromeleon management system, Dionex) and comparison with standard curves.

### LC-MS analyses

Detection of metabolites and quantification was performed by LC-MS. 10 µL of samples were analyzed using UPLC–MS with UPLC (Acquity H-class, Waters) coupled to a single quadrupole mass spectrometer (SQD mass analyzer, Waters) using an C18 column (Acquity UPLC BEH C18 2.1 mm × 50 mm, 1.7 µm). Elution was performed at 40°C with a constant flow rate of 0.6 mL/min using a gradient of Acetonitrile (solvent B) and water (solvent A) both acidified with 0.1% formic acid as follows: starting at 15% B during 2 min, solvent B was then raised from 15% to 95% in 5 min and maintained at 95% up to 9.5 min before going back to initial conditions at 9.8 min during 3 minutes before next injection if needed. Compounds were detected in electrospray positive ion mode by setting SQD parameters as follows: source temperature 130°C; desolvation temperature 400°C, and nitrogen flow: 1000 L.h-1 with mass range from m/z 800 to 1550. Surfactins were quantified based on their retention times and masses compared to commercial standards (98% purity, Lipofabrik).

### Induction of systemic resistance and ROS measurements

ISR assays were performed as previously described (39) on 4 weeks-old tobacco plants cultivated in hydroponic conditions using Hoagland solution as nutrient base. Plants were treated with pure surfactin at the root level and infected on leaves by applying a spore suspension of the phytopathogen *Botrytis cinerea* prepared as detailed previously (39). Spreading lesions occurred starting from 48h post-infection and the diameter size was measured two days later. Five plants were used per treatment and experiments were repeated independently at least twice. For determination of cytoplasmic ROS stimulation, fluorescent probe (DCFH-DA) was used. Plants used in this experiment were grown on Hoagland medium for two weeks as described above. Experiments were performed on nine samples per treatment each containing three root segments (approx 100 mg FW) collected from different plants (n=9). Roots were treated with 50 µM DCFH-DA for 10 minutes, rinsed with PBS upon removing the probe, and finally treated. All the operations were conducted in a 96-well black microplate. Fluorescence measurements were performed on a Spark (Tecan) microplate reader (exc 485 nm; em = 535 nm) with readings every 10 minutes. Stimulation of apoplastic hydrogen peroxide production in root cells was measured via chemiluminescence (ferricyanide-catalysed oxidation of luminol). Means and standard deviations were calculated from measurements performed on three samples per treatment, each containing three root segments (approximatively 100 mg FW) collected from different plants. Extracellular ROS in tomato roots was conducted according to Bisceglia *et al*. (80) with minor changes. Namely, instead of leaf discs, tomato roots, three segments (approximatively 100 mg FW from the same plant) per sample, were used. Plants were grown for two weeks on Hoagland medium, and chemiluminescence was measured in Tecan Spark plate reader.

### ITC analysis

ITC analyses were performed with a VP-ITC Microcalorimeter (Microcal). The calorimeter cell (volume of 1.4565 mL) was filled with a 10 µM (below the CMC concentration) surfactin solution in buffer (Tris 10mM, NaCl 150mM, 1mM EDTA at pH 8.5). The syringe was filled with a suspension of LUV at a lipid concentration of 5 mM. A series of 10µl injections was performed at constant time intervals (6 min) at 25°C. The solution in the titration cell was stirred at 305 RPM. Prior to each analysis, all solutions were degassed using a sonicator bath. The heats of dilution of vesicles were determined by injecting vesicles in buffer and subtracted from the heats determined in the experiments. Data were processed by software Origin 7 (Originlab) using the cumulative model described by Heerklotz and Seelig (81). All measurements were repeated at least three times with two different vesicle preparations.

### Leakage assays

Membrane permeabilization was followed as described by Van Bambeke *et al*. (82). Release of 8-hydroxypyrene-1,3,6 trisulfonic acid (HTPS) coentrapped with and quenched by p-xylene-bis-pyridinium bromide (DPX) from liposomes can be monitored by the fluorescence increase upon dilution following their leakage from the vesicles. Surfactin C12 or Surfactin C14 was added from a stock solution in DMSO and fluorescence intensities were immediately recorded. The percentage of HPTS released was defined as [(Ft-Fcontr)/(Ftot-Fcontr)] /100, where Ft is the fluorescence signal measured after 15 min in the presence of Surfactin C12 or Surfactin C14, Fcontr is the fluorescence signal measured at the same time for control liposomes, and Ftot is the total fluorescence signal obtained after complete disruption of the liposomes by 0.05% Triton X-100. All fluorescence determinations were performed at room temperature on a Perkin Elmer LS-50B Fluorescence Spectrophotometer (Perkin-Elmer Ltd.) using λexc of 450 nm and a λem of 512 nm.

### Statistical analyses

All statistical analyses were performed on GraphPad prism. Before each statistical analysis, variance homoscedasticity was verified by using a Brown-Forsythe test. ANOVA analysis was used for multiple comparison and significant differences were indicate by different letters. Statistical differences between means were evaluated by two-tailed Student’s t-test. Number or biological replicates used for each experiment are indicated in the corresponding figure legend. P-Values are indicated in the figure legends.

## Supporting information

Supplementary discussion

## Data availability

The RNA-seq datasets produced for this study are deposited at https://www.ebi.ac.uk/ena/ under the project reference PRJEB39762. All other datasets analyzed for this study are included in the supplementary files. The Genome Resulting assembly of the GA1 strain was deposited in the GenBank database under the accession numbers CP046386 and CP046387.

## Acknowledgments

This work was supported by the EU Interreg V France-Wallonie-Vlaanderen portfolio SmartBiocontrol (Bioprotect and Bioscreen projects, avec le soutien du Fonds européen de développement régional - Met steun van het Europees Fonds voor Regionale Ontwikkeling), by the PDR research project ID 26084552 from the F.R.S.-FNRS (National fund for Scientific Research in Belgium) and by the EOS project ID 30650620 from the FWO/F.R.S.-FNRS. FB is recipient of a F.R.I.A. fellowship (Formation à la Recherche dans l’Industrie et l’Agriculture) and MO is senior research associate at the F.R.S.- F.N.R.S. We are grateful to the KU Leuven HPC infrastructure and the Flemish Supercomputer Center (VSC) for providing the computational resources and services to perform the RNA-seq analysis. We gratefully acknowledge Claire Bertrand and Loïc Ongena for critically reading the manuscript.

## Notes

### Competing Interest Statement

The authors have declared no competing interest.

