## Supplementary material for "Surfactin stimulated by pectin molecular patterns and root exudates acts as a key driver of *Bacillus*-plant mutualistic interaction": Fig S6

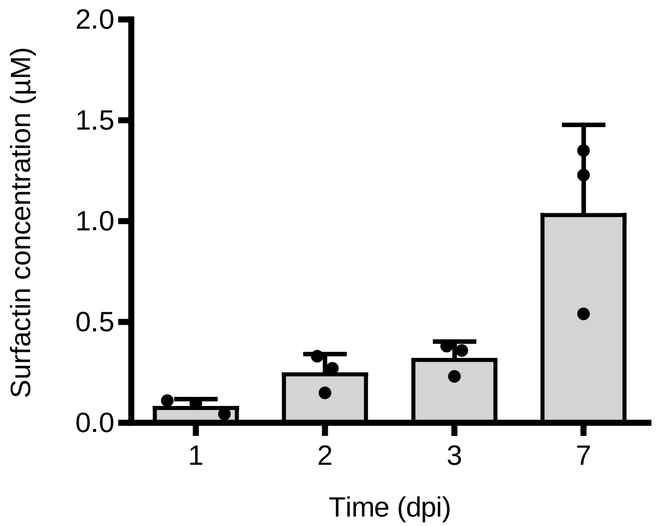


**Fig. S7:** Estimated surfactin concentration surrounding the rhizoplan in a solid matrix. Surfactin was quantified based on the amounts measured by UPLC-MS in three extracts (mean +- sd), each prepared from 3 roots and surrounding gelified medium from 3 individual plantlets. Surfactin concentration (µM) was calculated based on the mean value.
