## Supplementary material for "Surfactin stimulated by pectin molecular patterns and root exudates acts as a key driver of *Bacillus*-plant mutualistic interaction": Fig S5

a

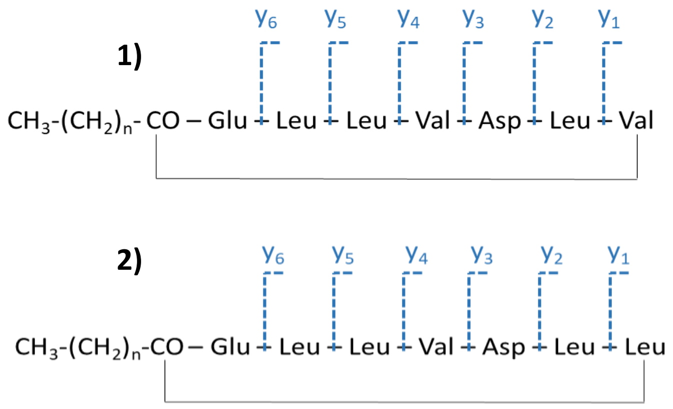


b

| Fragment Ion ID | Fragment ion Formula | Fragment m/z | Detected Mass Error (ppm) |
|---|---|---|---|
| **C13 Leu$^7$ surfactin (precursor ion : m/z = 1008,659112)** | | | |
| **y6** | C33H61N6O9 | 685,449454 | -0,3 |
| **y5** | C27H50N5O8 | 572,36539 | -1,7 |
| **y4** | C21H39N4O7 | 459,281326 | 0,1 |
| **y3** | C16H30N3O6 | 360,212912 | -2,5 |
| **y2** | C12H25N2O3 | 245,185969 | -1,1 |
| **y1** | C6H14NO2 | 132,101905 | -0,2 |
| **C13 Val$^7$ surfactin (precursor ion : m/z = 994,643462)** | | | |
| **y6** | C32H59N6O9 | 671,433804 | 1,2 |
| **y5** | C26H48N5O8 | 558,34974 | #N/A |
| **y4** | C20H3 N4O7 | 445,265676 | -2,7 |
| **y3** | C15H28N3O6 | 346,197262 | -1,1 |
| **y2** | C11H23N2O3 | 231,170319 | 0,2 |
| **y1** | C5H12NO2 | 118,086255 | 1,4 |
| **C14 Leu$^7$ surfactin (precursor ion : m/z = 1022,674762)** | | | |
| **y6** | C33H61N6O9 | 685,449454 | 0,3 |
| **y5** | C27H50N5O8 | 572,36539 | -1,9 |
| **y4** | C21H39N4O7 | 459,281326 | 1,5 |
| **y3** | C16H30N3O6 | 360,212912 | 0,9 |
| **y2** | C12H25N2O3 | 245,185969 | 0,7 |
| **y1** | C6H14NO2 | 132,101905 | 1,5 |
| **C14 Val$^7$ surfactin (precursor ion : m/z = 1008,659112)** | | | |
| **y6** | C32H59N6O9 | 671,433804 | 0,2 |
| **y5** | C26H48N5O8 | 558,34974 | -0,8 |
| **y4** | C20H37N4O7 | 445,265676 | 1,9 |
| **y3** | C15H28N3O6 | 346,197262 | 1,5 |
| **y2** | C11H23N2O3 | 231,170319 | -1,5 |
| **y1** | C5H12NO2 | 118,086255 | 1,7 |
| **C15 Leu$^7$ surfactin (precursor ion : m/z = 1036,690412)** | | | |
| **y6** | C33H61N6O9 | 685,449454 | -0,3 |
| **y5** | C27H50N5O8 | 572,36539 | -0,9 |
| **y4** | C21H39N4O7 | 459,281326 | -0,9 |
| **y3** | C16H30N3O6 | 360,212912 | -1,5 |
| **y2** | C12H25N2O3 | 245,185969 | -1,3 |
| **y1** | C6H14NO2 | 132,101905 | 0,9 |
| **C15 Val$^7$ surfactin (precursor ion : m/z = 1022,675762)** | | | |
| **y6** | C32H59N6O9 | 671,433804 | 0,6 |
| **y5** | C26H48N5O8 | 558,34974 | -0,6 |
| **y4** | C20H3 N4O7 | 445,265676 | -0,3 |
| **y3** | C15H28N3O6 | 346,197262 | 0,9 |
| **y2** | C11H23N2O3 | 231,170319 | #N/A |
| **y1** | C5H12NO2 | 118,086255 | 3,4 |

c

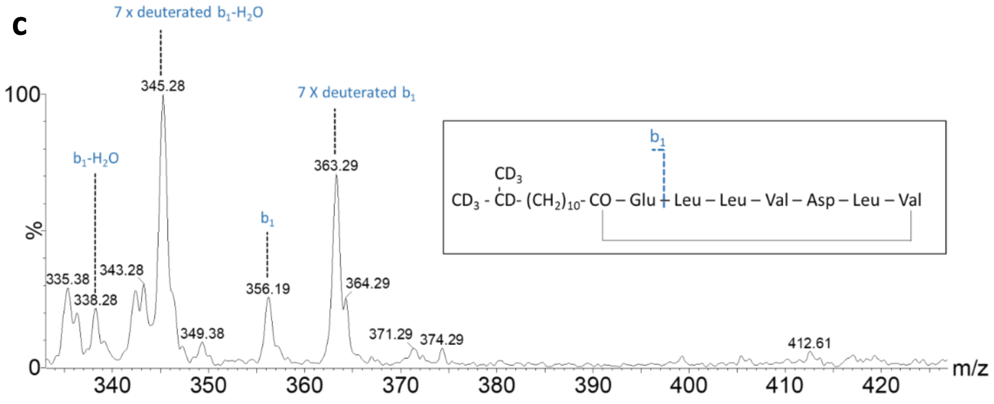


**Fig. S5: Surfactome variability: ab** HR-MSMS analyses. **a** Schematic representation of surfactin fragmentation. 1) surfactin Leu$_7$; 2) surfactin Val$_7$ **b** List and mass error of detected y-ions after fragmentation of surfactins produced in EM for C13 to C15 Leu$^7$ and Val$^7$ surfactins. Impact of medium supplementation with deuterated L-Val-d$^8$ on *B. velezensis* surfactome. Precursor feeding with 8 time deuterated valine will result in a mass increment of 7 mass unit in iso-even fatty acid (i.e. iso-C14; see insert) due the loss of α-deuterium during the transamination step [1]. Fragmentation of iso-C14 surfactin show a high proportion of deuterated b1 and b1-$H_2O$ fragment (m/z = 363 and 345 respectively). A small proportion of non-deuterated b1 and b1-$H_2O$ is also visible in the spectrum (m/z = 356 and m/z = 338 respectively).
