## Supplementary material for "Surfactin stimulated by pectin molecular patterns and root exudates acts as a key driver of *Bacillus*-plant mutualistic interaction": Fig S4

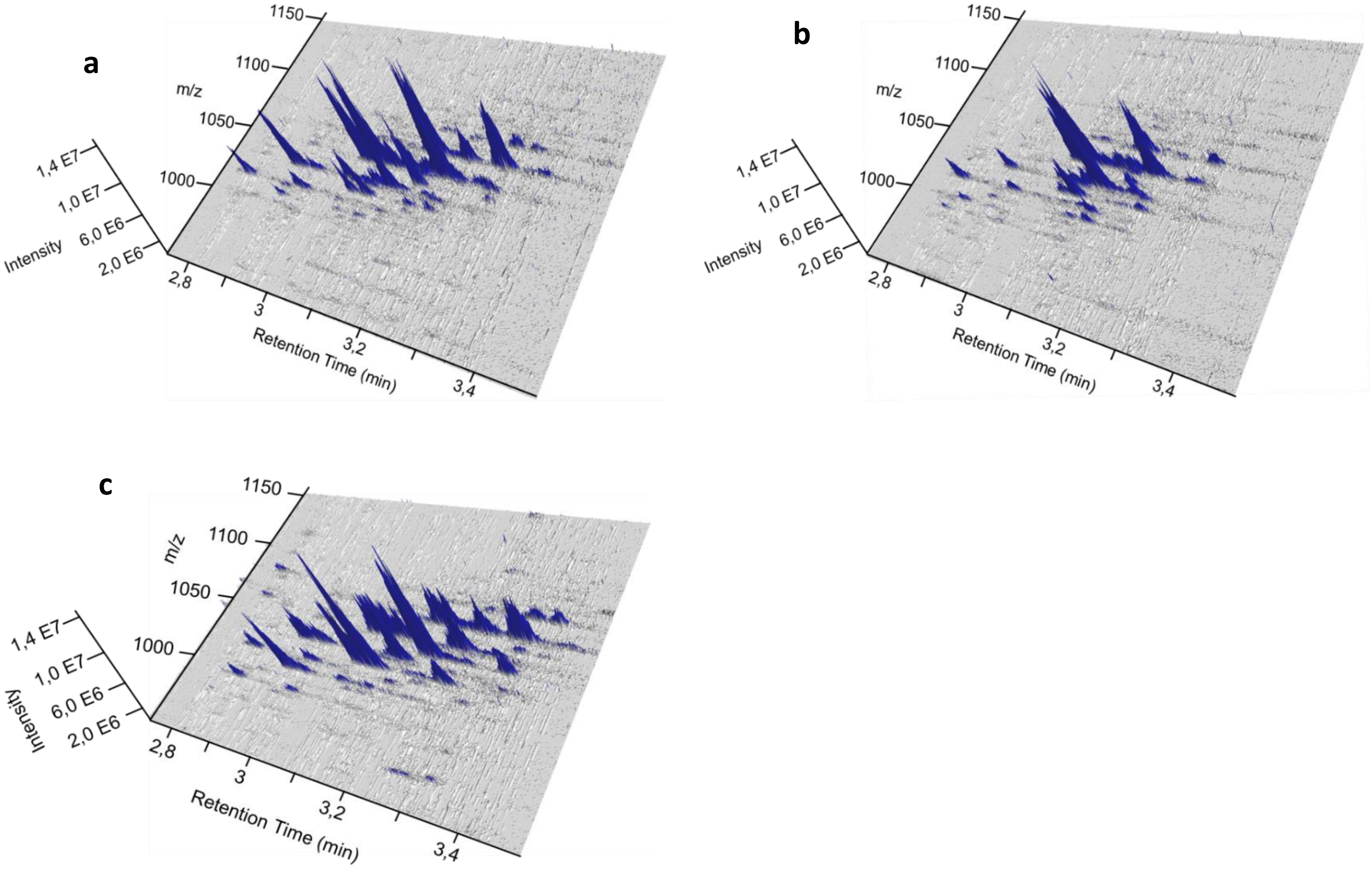


**Fig. S4: UPLC-MS 3D representation of *B. velezensis* surfactin pattern diversity produced in REM medium (a), in natural root exudates (b), or in planta (c).** The X axis indicates the retention time (min), the Y axis the mass to charge ratio (m/z) and the Z axis the peak intensity (AU). Each blue peak represents a surfactin homologue.
