## Supplementary material for "Surfactin stimulated by pectin molecular patterns and root exudates acts as a key driver of *Bacillus*-plant mutualistic interaction": Fig S3

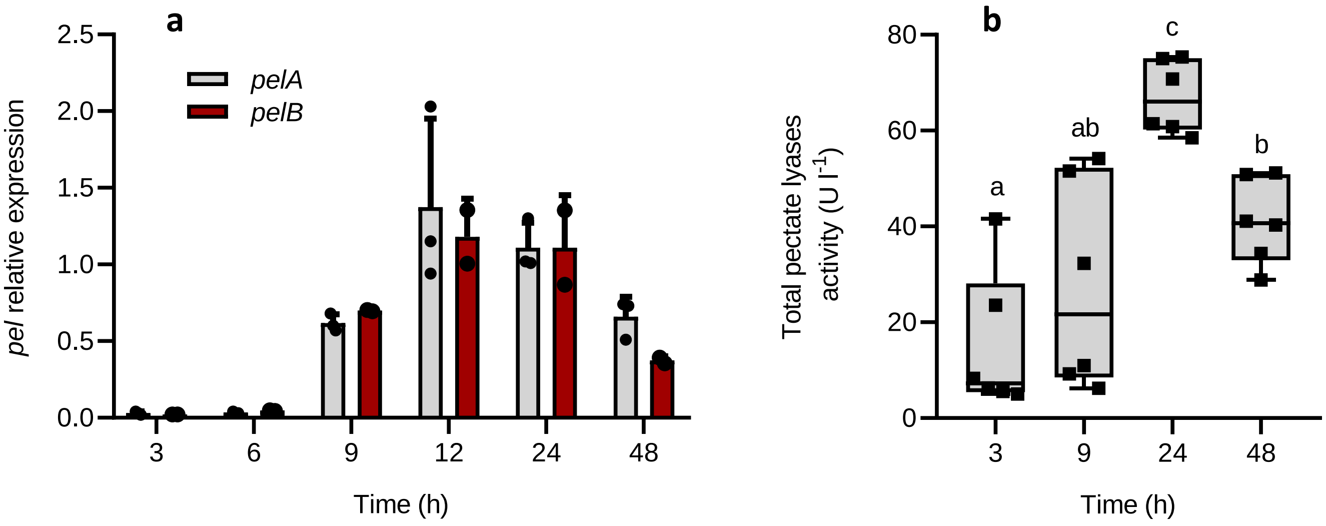


**Fig. S3: Characterization of *pel* expression and pectate lyase activity in GA1. a** Evolution of *pelA* (grey) and *pelB* (red) expression pattern (n=3) For each time point, means ± std err. from three biological replicates of one experiment are shown **b** Evolution of global pectate lyase activity in a 48h time course experiment. The box plots encompass the 1st and 3rd quartile, the whiskers extend to the minimum and maximum points, and the midline indicates the median (n=6 biological replicates of two experiments). Significate differences are indicated by different letters (n=6).
