## Supplementary material for "Surfactin stimulated by pectin molecular patterns and root exudates acts as a key driver of *Bacillus*-plant mutualistic interaction": Fig S2

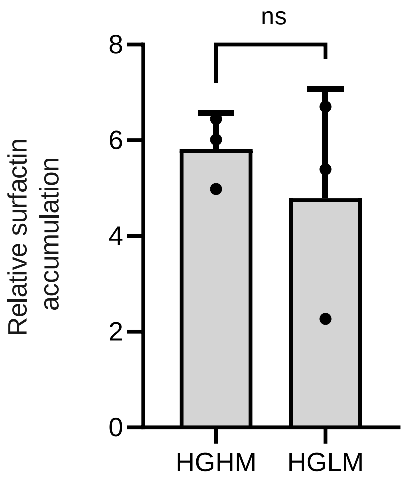


**Fig. S2: Relative surfactin accumulation by GA1 cells at early growth phase ($OD_{600}$=0.2) after addition of low (HGLM) or high (HGHM) methyl-esterified HG**. Means ± std err. from three biological replicates of one experiment are shown ns= non significate.
