## Supplementary material for "Surfactin stimulated by pectin molecular patterns and root exudates acts as a key driver of *Bacillus*-plant mutualistic interaction": Fig S1

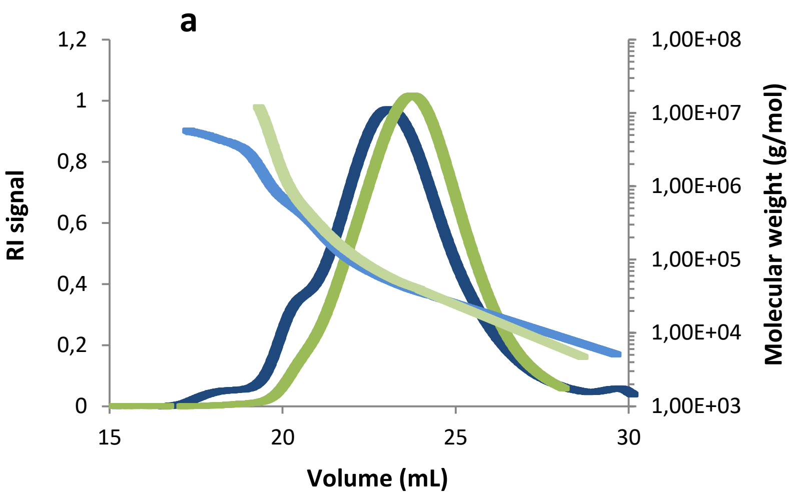


**b**

| | HGLM | HGHM |
|---|---|---|
| **Mn (g/Mol)** | $3.328 \times 10^4$ | $3.693 \times 10^4$ |
| **MW (g/mol)** | $9.823 \times 10^4$ | $1.947 \times 10^5$ |
| **Polydispersity (Mw/Mn)** | 3.030 | 5.273 |

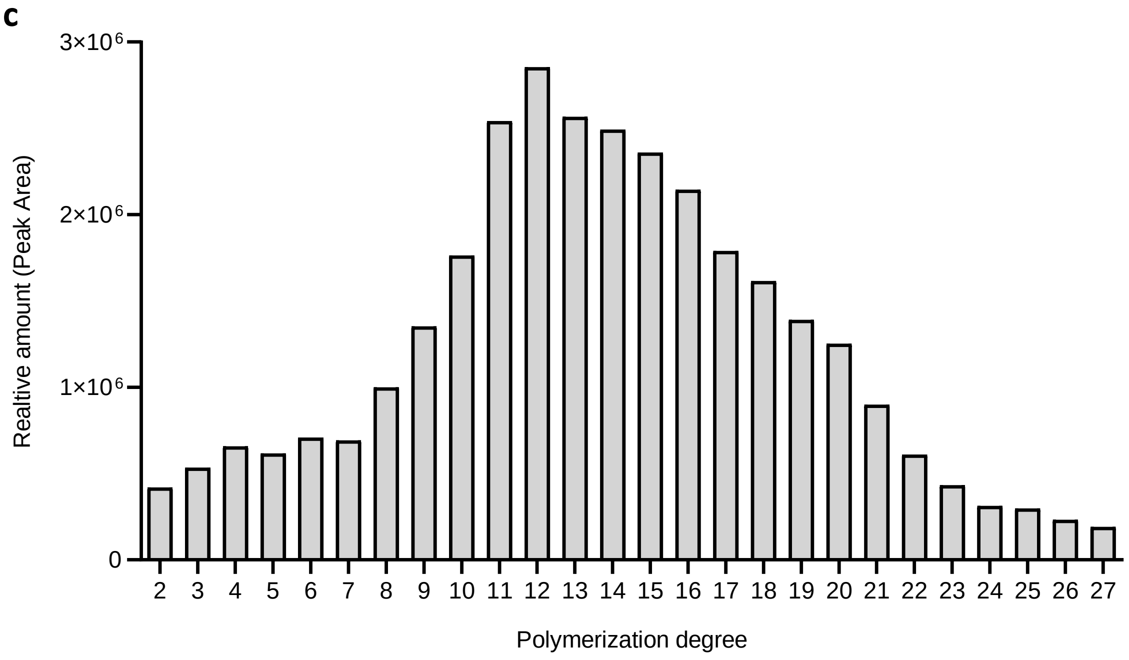


**Fig. S1: ab** SEC-Malls characterization of high and low methylated homogalacturonan **a** SEC-Malls profile of high (blue) and low (green) methylated homogalacturonan. Light curves represent the molecular weight distribution, dark curves represent the RI signal. **b** SEC-Malls results. Mn: number average molecular weight, Mw : weight-average molecular weight, and polydispersity values (Mw/Mn). **c** Characterization of oligogalacturonides (OG) polymerization degree by HILIC-QTOF
