## Supplementary discussion for "Surfactin stimulated by pectin molecular patterns and root exudates acts as a key driver of *Bacillus*-plant mutualistic interaction"

Surfactin producing strains are always producing the cLP as a mixture of different structural variants. Specific amino acids substitution in the peptide as well as differences in the fatty acid (FA) chain length and branching type represent the main structural variations found in this mixture of surfactins. Two main factors are found to be involved in this diversity.

On one hand, the availability of precursors as well as the balance between these latter is an important factor that drive the surfactin structural diversity. Indeed, feeding the bacteria with specific amino acids will result in changes in the ratio between the different variants (peptidic and lipidic) as well as production of new variants that are not produced in normal lab conditions (1, 2). Among the important amino acids for surfactin production, isoleucine, valine and leucine (branched chain amino acids or BCAAs) constitute the major precursors (5 out of 7). The key role of BCAAs is even more important since the branched chain fatty acids (BCFAs) tail of surfactin biosynthetic pathway also require these BCAAs as precursors for their biosynthesis. Indeed, Ile, Leu and Val are precursors of odd anteiso-FAs, odd iso-FAs and even iso FAs, respectively (3). This makes thus from the BCAAs balance the key factor for most part of the building blocks of surfactin. Therefore, deciphering the BCAAs biosynthetic pathways as well as its regulation can bring some clues on what can be involved in the balance changes of surfactin precursors. Even if the biosynthetic pathways responsible for these three amino acids share part of the enzyme coding genes (*ilvBCDE*), precursors as well as some additional genes are specifically involved in the biosynthesis of each BCAA. Ile require Thr and pyruvate to be synthesized via the action of enzymes encoded by *ilvABCDE.* Val and Leu share the first part of the transformation of pyruvate into α-ketoisovalerate thanks to enzymes encoded by *ilvBCD* genes. This intermediate can be then transformed into Valine via IlvE enzyme or can further react with acetyl-CoA and undergo enzymatic transformation involving *leuABCD* and *ilvE* genes to produce Leucine (4). Regulation of these enzymes is most probably also an important factor to understand the balance changes that can occur in the precursor's pools. The pleiotropic regulator has been shown to monitor and regulate the intracellular concentrations of BCAAs. Indeed, BCAAs (as well as GTPs) can bind to specific pockets of CodY tetramers and induce structural changes of the regulator that will expose the DNA binding domain of the regulator (5). By binding to specific DNA sequences, CodY will play a role in the regulation of hundreds genes, mainly as a repressor, including ilv and leu genes (6, 7). Interestingly, in B. subtilis, derepression of CodY will result only in a slight increase of Leu and Ile intracellular concentrations (2-fold) but up to 6-fold higher Val concentration as well as 24-fold increase expression of *ybgE* coding for an enzyme involved in BCFAs biosynthesis (7). The reasons of this differential impact of the regulator even if the enzymes are shared in the biosynthesis of the three amino acids is still unknown. It has been hypothesized that Leu and Ile can be excreted outside of the cells, or that some CodY-independent regulation is involved in Leu/Ile regulation.

On the other hand, such a diversity in the structure of surfactins produced by a same strain will never be possible by a ribosomal mode of biosynthesis. The non-ribosomal machinery that is involved in the selection, activation and elongation of the nascent lipopeptides is indeed able to add different type of fatty acid as well as amino acids at certain position of the molecule. The mega enzymatic complex responsible for surfactin production is mainly composed of 7 seven modules (one per amino acid) that are subdivided into at least 3 specific domains: adenylation (A-domain), condensation (C domain) and PCP (peptidyl-carrier-protein domain) (8). Among those, A-domain is known to be responsible for amino acid selectivity. Indeed, each A-domain show a specific 8 amino acid binding pocket that is able to interact with a specific amino acid that will be activated and inserted into the peptide via the action of the C domain (9–12). It appears that some binding pockets of certain modules can show differential affinity for more than one amino acid and therefore can lead to the production of peptidic variants. Similarly, it appears that addition of fatty acid occurring at the beginning of the surfactin biosynthesis is under the specific selectivity of first C-domain of surfactin operon  (13).

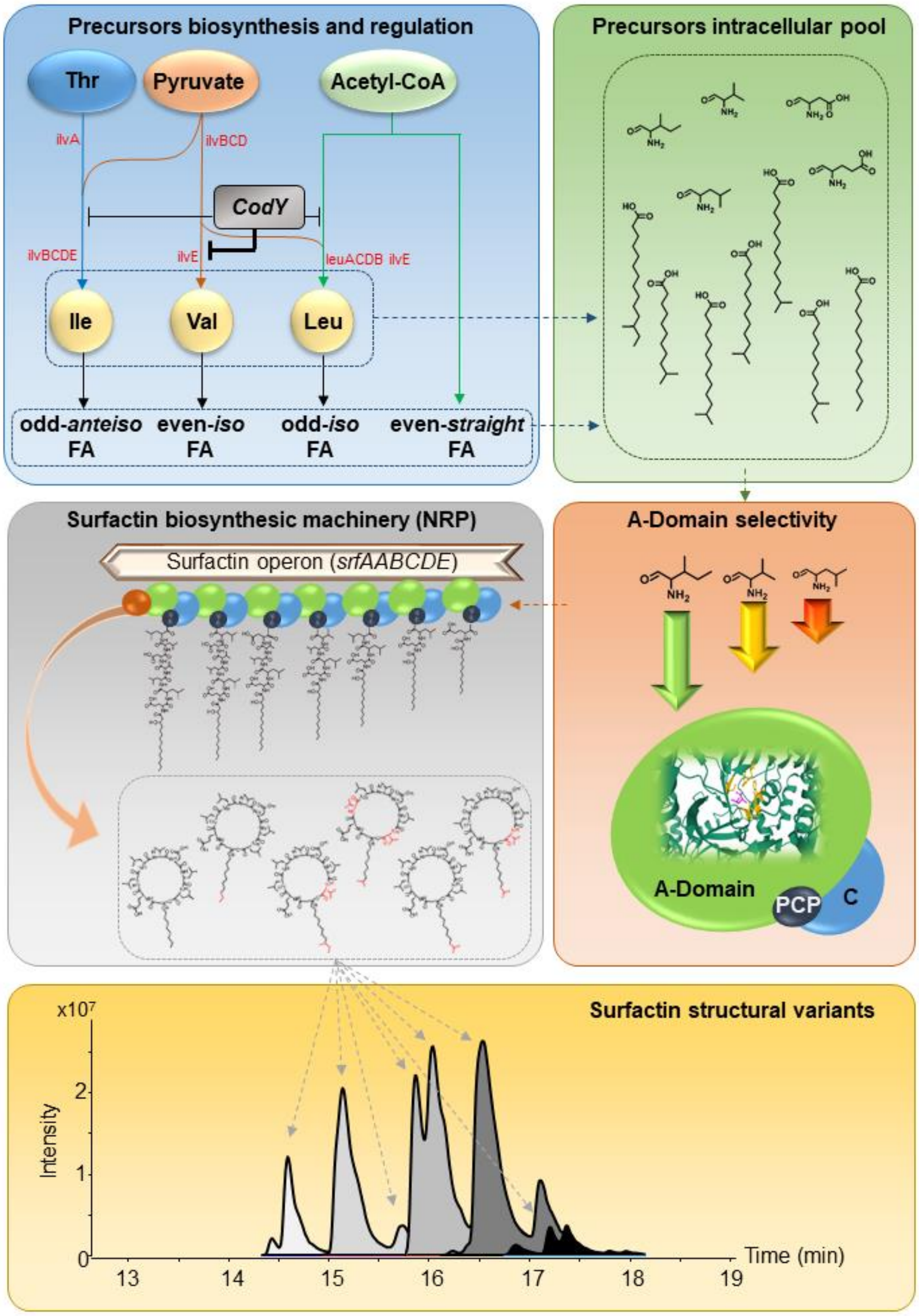


**Supplementary discussion Fig. 1:** Non ribosomal biosynthesis via NRP synthetases and regulation by pleiotropic transcription factors such as CodY both drive diversity and ratio of surfactin's precursors in the intracellular pool (i.e. branched chain amino acids and fatty acids, blue panel). In turn, relative amounts in the intracellular pool (green panel) as well as the selectivity of each adenylation domain regarding the type of amino acid it activates (orange panel) lead to the production by the NRP machinery of different surfactin variants (grey panel) detected in reversed phase LC-MS (yellow panel).
