## Supplementary material for "Surfactin stimulated by pectin molecular patterns and root exudates acts as a key driver of *Bacillus*-plant mutualistic interaction"

**Quantitative RT-PCR.** RT-qPCR was undertaken in a StepOnePlus™ Real-Time PCR System (Applied Biosystems) apparatus in a 20 µL reaction mixture containing respectively 10 µL of Luna Universal One-Step Reaction Mix (New England BioLabs), 1 µL of Luna WarmStart® RT Enzyme Mix, 0.4 µM of each primer and 100 ng of RNA. The reaction conditions were as follow: a 10 min reverse transcription step at 55°C, a 1 min initial denaturation step at 95°C and 40 cycles of 10 s at 95°C and 30 s at 60°C. Each cycle was achieved by a fluorescence plate read. In order to check the absence of secondary products, melting curves were realized from 65 to 95°C with an increase rate of 0.5°C/5 s. The transcription level of interest genes was analyzed using the mathematical model proposed by Pfaffl (1). The *gyrA* gene encoding DNA gyrase subunit A was used as the reference gene for the normalization of gene expression level with a constitutively expressed gene.

**Size Exclusion Chromatography (SEC-MALLS).** $NaNO_3$, 0.1 M and $NaN_3$, 2.5mM, used as carrier, was filtered through a 0.02 µm, 47 mm membrane filter (Anotop 47, Whatman), carefully degassed. Samples (5 mg/mL) were filtered through a 0.45 mm membrane filter (Grace Altech) and were injected through a 100 µL full loop. The SEC line consisted of an SB-G guard column as protection and three columns in series (SB-806 HQ, SB-804 HQ and SB-803 HQ, 300 mm L × _8 mm I.D., Shodex Showa Denko K.K.). The elution was performed with a flow rate of 0.5 $mL.min^{-1}$ (LC-20AD, Shimadzu). Detection was achieved with a light scattering detector (MiniDAWN TREOS II, Wyatt Technology Corporation) and a refractive index detector (RID-10 A, Shimadzu). Data acquisition was performed using ASTRA 7.2.2 software.

**Pectate lyase activity measurement.** Total pectate lyase activity of culture supernatants was measured by spectrophotometry. Accumulation of unsaturated oligogalacturonides generated by the enzymes via the mechanism of β-elimination was evaluated by measuring the increase of absorbance at 232 nm. The molar extinction coefficient for unsaturated oligogalacturonides at 232 nm is 4600 $M^{-1}$ $cm^{-1}$ (2). Briefly, 62.5 µl of filtered culture supernatant were preheated at 25°C and added to 437.5 µL of HG (5g/l) dissolved in 0.057 M phosphate buffer at pH 8. The reaction mixture was then incubated in a Thermomixer at 40°C, 200 rpm, during 180 minutes. Sampling was performed after 10, 30, 60, 120, 180 min and the enzymatic reaction was stopped by placing the samples into ice during 10 min. One unit of enzymatic activity (U) was considered as equivalent of 1 µmol of unsaturated oligogalacturonides produced per min.

**Surfactin structural elucidation.** Structural elucidation of surfactin was performed by ultra-high-performance liquid chromatography coupled to high resolution tandem mass spectrometry with a heated electrospray ionization source as interface (UPLC-HESI-HRMS/MS). Chromatographic separation of analytes was performed on an Acquity UPLC I-Class Chromatographic system (Waters) by injection of 10 µL of extracts on an Acquity UPLC BEH C18 column (1.7 µm, 2.1x50 mm) column (Waters). Elution was performed at 40 °C with a constant flow rate of 0.6 mL/min using a 7 min gradient of ACN (A): 0.1 % HFo (B) as follows: 1 min at 30 % A, from 30 % A to 95 % A in 3.6 min and maintained 1 min at 95 % A. Accurate mass measurements were performed on a Quadrupole-Orbitrap instrument (Q-Exactive Plus, Thermo Electron GmbH) working in full scan plus data dependent (dd-MS2) and parallel reaction monitoring (PRM) acquisition modes equipped with an HESI probe as interface working in positive ionization mode (HESI(+)). LC eluent was electrosprayed in the HESI interface at the following conditions: spray voltage 3.0 kV, capillary temperature 400 °C, sheath gas 60 a.u., auxiliary gas 20 a. u., maximum spray current 100 µA, probe heater temperature 100 ºC, and S-Lens RF Level 50. Full scan mass spectra (MS) were acquired in the mass range from 400 to 1500 m/z at a mass resolution of 140,000 full width at half maximum (FWHM) (m/z 400), automatic gain control (AGC) target of 3x106 or maximum injection time of 200 ms. For dd-MS2 mode, TopN-MS/MS was set to 5. Mass tandem spectra (MS2) were acquired in the mass range from 200 to 2000 m/z at a mass resolution of 17,500 FWHM (m/z 400), automatic gain control (AGC) target of 2x105 or maximum injection time of 150 ms, isolation window of 2 m/z and stepped normalized collision energy (NCE) of 21.2, 25 and 28 eV. HRMS of proton, sodium and potassium adduct and HRMS2 of protonated molecular ion, as well as isotopic distributions and chromatographic retention time, were used for confirmation. Accurate mass measurements were performed on the marker molecular ion and fragments, characteristic of every isomer, with mass accuracy below 2.5 ppm. Data were acquired and processed using Xcalibur software version 3.1. Mass calibration was performed with a Thermo Scientific™ Pierce™ LTQ ESI Positive Ion Calibration Solution, a mixture of highly purified ionizable molecules specifically designed for positive mode calibration of Orbitrap instruments.
