## Supplementary table 2 for "Surfactin stimulated by pectin molecular patterns and root exudates acts as a key driver of *Bacillus*-plant mutualistic interaction"

**Supplementary table 2:** Mapping metrics. Absolute number and percentage of reads mapped to the annotated reference genome.

| Sample | Total number of trimmed reads | Mapped reads | [%] |
|---|---|---|---|
| GA1_2h30_R1 | 10153078 | 9791294 | 96.44% |
| GA1_2h30_R2 | 11793634 | 11407196 | 96.72% |
| GA1_2h30_R3 | 12515446 | 11942542 | 95.42% |
| GLM_2h30_R1 | 11424956 | 11002586 | 96.30% |
| GLM_2h30_R2 | 11108682 | 10646558 | 95.84% |
| GLM_2h30_R3 | 13085518 | 12483206 | 95.40% |
| GA1_5h_R1 | 13425432 | 12994028 | 96.79% |
| GA1_5h_R2 | 13946330 | 13523602 | 96.97% |
| GA1_5h_R3 | 11718380 | 11361472 | 96.95% |
| GLM_5h_R1 | 11913410 | 11534496 | 96.82% |
| GLM_5h_R2 | 13400252 | 12997016 | 96.99% |
| GLM_5h_R3 | 12352654 | 11926200 | 96.55% |
| GA1_8h_R1 | 10770746 | 10366778 | 96.25% |
| GA1_8h_R2 | 11264570 | 10894940 | 96.72% |
| GA1_8h_R3 | 13098010 | 12673498 | 96.76% |
| GLM_8h_R1 | 13468736 | 13040536 | 96.82% |
| GLM_8h_R2 | 11912620 | 11547116 | 96.93% |
| GLM_8h_R3 | 13169894 | 12737898 | 96.72% |
