## Supplementary table 1 for "Surfactin stimulated by pectin molecular patterns and root exudates acts as a key driver of *Bacillus*-plant mutualistic interaction"

**Supplementary table 1:** Conservation of the pectate lyases *(*pel) genes in the *"Operational Group B.* amyloliquefaciens"

| | Genome size (Mb) | PL1 | | | |
|---|---|---|---|---|---|
| | | Pectate lyase | | | |
| ***Bacillus velezensis* GA1** | | GL331_08735 | | GL331_04125 | |
| | | *pelA* | | *pelB* | |
| | 3.86782 | Cov (%) | Id (%) | Cov (%) | Id (%) |
| *Bacillus amyloliquefaciens* DSM7 | 3.9802 | 100 | 93.52 | 100 | 95.02 |
| *Bacillus amyloliquefaciens* HK1 | 4.00284 | 100 | 93.52 | 100 | 95.02 |
| *Bacillus amyloliquefaciens* LL3 | 4.00199 | 100 | 93.92 | 100 | 94.93 |
| *Bacillus amyloliquefaciens* MT45 | 3.89752 | 100 | 93.84 | 100 | 95.49 |
| *Bacillus amyloliquefaciens* RD7-7 | 3.68821 | 100 | 93.52 | 100 | 95.96 |
| *Bacillus amyloliquefaciens* SRCM101267 | 4.08824 | 100 | 93.52 | 100 | 95.02 |
| *Bacillus amyloliquefaciens* TA208 | 3.93751 | 100 | 93.92 | 100 | 95.02 |
| *Bacillus amyloliquefaciens* XH7 | 3.9392 | 100 | 93.92 | 100 | 95.02 |
| *Bacillus amyloliquefaciens* YP6 | 4.00962 | 100 | 94.23 | 100 | 95.4 |
| *Bacillus siamensis* SCSIO 05746 | 4.28071 | 100 | 94.39 | 100 | 95.77 |
| *Bacillus velezensis* 10075 | 4.33983 | 100 | 98.1 | 100 | 97.84 |
| *Bacillus velezensis* 131-4 | 3.89303 | 100 | 99.61 | 100 | 99.15 |
| *Bacillus velezensis* 157 | 4.02069 | 100 | 97.79 | 100 | 97.93 |
| *Bacillus velezensis* 1B-23 | 4.14106 | 100 | 97.79 | 100 | 97.84 |
| *Bacillus velezensis* 8-2 | 3.89307 | 100 | 99.61 | 100 | 99.15 |
| *Bacillus velezensis* 83 | 3.9979 | 99 | 98.02 | 100 | 97.75 |
| *Bacillus velezensis* 9912D | 4.24158 | 100 | 98.1 | 100 | 97.65 |
| *Bacillus velezensis* 9D-6 | 3.96373 | 100 | 97.39 | 100 | 98.03 |
| *Bacillus velezensis* AL7 | 3.99598 | 100 | 97.87 | 100 | 98.5 |
| *Bacillus velezensis* ALB65 | 4.04167 | 100 | 97.95 | 100 | 98.5 |
| *Bacillus velezensis* ALB69 | 4.04661 | 100 | 97.95 | 100 | 98.87 |
| *Bacillus velezensis* ALB79 | 3.98291 | 100 | 97.31 | 100 | 97.93 |
| *Bacillus velezensis* ANSB01E | 3.92984 | 100 | 97.79 | 100 | 97.56 |
| *Bacillus velezensis* AP183 | 4.00644 | 100 | 97.08 | 100 | 98.12 |
| *Bacillus velezensis* ARP23 | 4.01887 | 100 | 97.71 | 100 | 97.93 |
| *Bacillus velezensis* AS43.3 | 3.96137 | 100 | 97.31 | 100 | 97.84 |
| *Bacillus velezensis* At1 | 3.88899 | 100 | 97.71 | 100 | 98.22 |
| *Bacillus velezensis* ATR2 | 4.00675 | 100 | 98.1 | 100 | 97.84 |
| *Bacillus velezensis* B15 | 4.00675 | 100 | 97.55 | 100 | 98.12 |
| *Bacillus velezensis* B25 | 3.86276 | 100 | 100 | 100 | 100 |
| *Bacillus velezensis* B4 | 3.9198 | 100 | 99.21 | 100 | 98.69 |
| *Bacillus velezensis* Bac57 | 4.2349 | 100 | 98.1 | 100 | 98.22 |
| *Bacillus velezensis* BCSo1 | 3.71352 | 100 | 97.31 | 100 | 98.31 |
| *Bacillus velezensis* BIM B-439D | 3.97895 | 100 | 97.47 | 100 | 98.22 |
| *Bacillus velezensis* BS-37 | 4.01389 | 100 | 97.08 | 100 | 98.03 |

| | | | | | |
|---|---|---|---|---|---|
| *Bacillus velezensis* BvL103 | 3.98454 | 100 | 99.13 | 100 | 99.44 |
| *Bacillus velezensis* CAU B946 | 4.01986 | 100 | 99.37 | 100 | 99.25 |
| *Bacillus velezensis* CBMB205 | 3.92975 | 100 | 97.79 | 100 | 97.56 |
| *Bacillus velezensis* CC09 | 4.16715 | 100 | 96.68 | 100 | 97.93 |
| *Bacillus velezensis* CC178 | 3.91683 | 100 | 97.79 | 100 | 97.84 |
| *Bacillus velezensis* CGMCC 11640 | 4.38568 | 100 | 96.68 | 100 | 97.93 |
| *Bacillus velezensis* CMT-6 | 3.92849 | 100 | 99.53 | 100 | 99.72 |
| *Bacillus velezensis* CN026 | 3.99581 | 99 | 97.94 | 100 | 97.65 |
| *Bacillus velezensis* DH8030 | 3.99398 | 100 | 99.68 | 100 | 100 |
| *Bacillus velezensis* DKU_NT_04 | 4.32819 | 100 | 98.03 | 100 | 97.93 |
| *Bacillus velezensis* DR-08 | 3.92979 | 100 | 97.79 | 100 | 97.56 |
| *Bacillus velezensis* DSYZ | 4.32146 | 100 | 96.68 | 100 | 97.93 |
| *Bacillus velezensis* FJAT-46737 | 3.99598 | 100 | 97.37 | 100 | 98.12 |
| *Bacillus velezensis* FJAT-52631 | 3.92978 | 100 | 97.79 | 100 | 97.56 |
| *Bacillus velezensis* FS1092 | 4.24093 | 100 | 97.31 | 100 | 97.93 |
| *Bacillus velezensis* FZB42 | 3.91859 | 100 | 97.79 | 100 | 97.84 |
| *Bacillus velezensis* G341 | 4.00975 | 100 | 97.79 | 100 | 98.22 |
| *Bacillus velezensis* GFP-2 | 3.97522 | 100 | 98.89 | 100 | 98.31 |
| *Bacillus velezensis* GH1-13 | 4.14361 | 100 | 98.89 | 100 | 99.25 |
| *Bacillus velezensis* GQJK49 | 3.92976 | 100 | 97.79 | 100 | 97.56 |
| *Bacillus velezensis* GYL4 | 3.97508 | 100 | 97.63 | 100 | 97.93 |
| *Bacillus velezensis* Hx05 | 3.91387 | 100 | 99.61 | 100 | 99.06 |
| *Bacillus velezensis* IT45 | 3.93687 | 100 | 99.53 | 100 | 98.78 |
| *Bacillus velezensis* J01 | 4.17993 | 100 | 99.53 | 100 | 99.72 |
| *Bacillus velezensis* J7-1 | 3.89307 | 100 | 99.61 | 100 | 99.15 |
| *Bacillus velezensis* JJ-D34 | 4.10595 | 100 | 99.37 | 100 | 99.25 |
| *Bacillus velezensis* JS25R | 4.01444 | 100 | 97.79 | 100 | 98.59 |
| *Bacillus velezensis* JT3-1 | 3.9298 | 100 | 97.79 | 100 | 97.56 |
| *Bacillus velezensis* JTYP2 | 3.92979 | 100 | 97.79 | 100 | 97.56 |
| *Bacillus velezensis* K26 | 4.04735 | 100 | 98.03 | 100 | 97.75 |
| *Bacillus velezensis* KC41 | 4.11876 | 100 | 98.1 | 100 | 98.03 |
| *Bacillus velezensis* KD1 | 3.92197 | 100 | 99.05 | 100 | 99.81 |
| *Bacillus velezensis* KHG19 | 3.95336 | 100 | 97.55 | 100 | 98.31 |
| *Bacillus velezensis* L1 | 4.09058 | 100 | 97.55 | 100 | 98.87 |
| *Bacillus velezensis* LABIM40 | 3.97231 | 100 | 97.16 | 100 | 98.22 |
| *Bacillus velezensis* LB002 | 4.07686 | 100 | 99.68 | 100 | 100 |
| *Bacillus velezensis* LC1 | 3.92978 | 100 | 97.79 | 100 | 97.56 |
| *Bacillus velezensis* LDO2 | 3.94727 | 100 | 97.79 | 100 | 97.56 |
| *Bacillus velezensis* LFB112 | 3.94275 | 100 | 99.68 | 100 | 100 |
| *Bacillus velezensis* LG37 | 3.92 | 100 | 97.79 | 100 | 97.56 |
| *Bacillus velezensis* L-H15 | 3.93329 | 100 | 99.29 | 100 | 100 |
| *Bacillus velezensis* LM2303 | 3.98939 | 100 | 99.61 | 100 | 99.06 |
| *Bacillus velezensis* LPL-K103 | 3.90302 | 100 | 97.63 | 100 | 98.78 |
| *Bacillus velezensis* L-S60 | 3.90597 | 100 | 99.29 | 100 | 100 |
| *Bacillus velezensis* LS69 | 3.91776 | 100 | 97.79 | 100 | 97.56 |
| *Bacillus velezensis* Lzh-a42 | 4.2466 | 99 | 98.02 | 100 | 98.12 |

| | | | | | |
|---|---|---|---|---|---|
| *Bacillus velezensis* M75 | 4.00745 | 100 | 99.68 | 100 | 99.91 |
| *Bacillus velezensis* MBE1283 | 3.97993 | 100 | 99.68 | 100 | 99.34 |
| *Bacillus velezensis* MH25 | 4.11847 | 100 | 97.63 | 100 | 98.31 |
| *Bacillus velezensis* NAU-B3 | 4.20461 | 100 | 97.79 | 100 | 98.59 |
| *Bacillus velezensis* NJAU-Z9 | 3.87256 | 100 | 99.61 | 100 | 99.15 |
| *Bacillus velezensis* NJN-6 | 4.05255 | 100 | 99.68 | 100 | 100 |
| *Bacillus velezensis* NKG-1 | 4.19722 | 100 | 97.87 | 100 | 98.03 |
| *Bacillus velezensis* NY12-2 | 4.07491 | 100 | 98.03 | 100 | 97.93 |
| *Bacillus velezensis* ONU 553 | 3.93456 | 100 | 97.79 | 100 | 97.84 |
| *Bacillus velezensis* OSY-GA1 | 4.01 | 100 | 99.53 | 100 | 99.72 |
| *Bacillus velezensis* P34 | 3.90046 | 100 | 97.31 | 100 | 98.59 |
| *Bacillus velezensis* QST713 | 4.23376 | 100 | 97.31 | 100 | 97.93 |
| *Bacillus velezensis* S141 | 3.97458 | 100 | 97.71 | 100 | 98.12 |
| *Bacillus velezensis* S3-1 | 3.92977 | 100 | 97.79 | 100 | 97.56 |
| *Bacillus velezensis* S499 | 3.93593 | 100 | 99.53 | 100 | 98.78 |
| *Bacillus velezensis* SCDB 291 | 4.16257 | 100 | 98.18 | 100 | 99.44 |
| *Bacillus velezensis* SCGB 1 | 4.08549 | 100 | 98.18 | 100 | 99.44 |
| *Bacillus velezensis* SCGB 574 | 3.98915 | 100 | 97.79 | 100 | 98.59 |
| *Bacillus velezensis* SGAir0473 | 4.18428 | 100 | 98.1 | 100 | 98.03 |
| *Bacillus velezensis* SH-B74 | 4.10382 | 100 | 97.71 | 100 | 98.22 |
| *Bacillus velezensis* SQR9 | 4.11702 | 100 | 97.24 | 100 | 99.06 |
| *Bacillus velezensis* SRCM100072 | 4.00176 | 100 | 97.95 | 100 | 98.03 |
| *Bacillus velezensis* SRCM101413 | 4.20919 | 100 | 98.03 | 100 | 97.84 |
| *Bacillus velezensis* SRCM103616 | 4.23511 | 100 | 98.03 | 100 | 97.84 |
| *Bacillus velezensis* SRCM103691 | 4.13953 | 100 | 98.03 | 100 | 97.84 |
| *Bacillus velezensis* SRCM103788 | 4.1441 | 100 | 98.03 | 100 | 97.84 |
| *Bacillus velezensis* sx01604 | 3.92652 | 100 | 97.79 | 100 | 97.56 |
| *Bacillus velezensis* SYBC H47 | 3.88443 | 100 | 98.97 | 100 | 98.5 |
| *Bacillus velezensis* SYP B637 | 3.91555 | 100 | 97.79 | 100 | 97.84 |
| *Bacillus velezensis* T20E-257 | 3.90007 | 100 | 99.61 | 100 | 99.15 |
| *Bacillus velezensis* TB1501 | 3.97938 | 100 | 97.63 | 100 | 98.87 |
| *Bacillus velezensis* TJ02 | 4.06155 | 100 | 97.63 | 100 | 98.22 |
| *Bacillus velezensis* TrigoCor1448 | 3.9579 | 100 | 97.55 | 100 | 98.03 |
| *Bacillus velezensis* UCMB5007 | 3.98332 | 100 | 97.55 | 100 | 98.31 |
| *Bacillus velezensis* UCMB5033 | 4.07117 | 100 | 97.31 | 100 | 98.31 |
| *Bacillus velezensis* UCMB5036 | 3.91032 | 100 | 97.71 | 100 | 98.03 |
| *Bacillus velezensis* UCMB5044 | 3.9833 | 100 | 97.55 | 100 | 98.31 |
| *Bacillus velezensis* UCMB5113 | 3.88953 | 100 | 97.71 | 100 | 98.22 |
| *Bacillus velezensis* UFLA258 | 3.94721 | 100 | 97.79 | 100 | 98.4 |
| *Bacillus velezensis* UMAF6614 | 4.00514 | 100 | 97.95 | 100 | 98.87 |
| *Bacillus velezensis* UMAF6639 | 4.03464 | 100 | 97.63 | 100 | 98.03 |
| *Bacillus velezensis* UTB96 | 3.71568 | 100 | 99.29 | 100 | 99.91 |
| *Bacillus velezensis* V167 | 3.90447 | 100 | 97.79 | 100 | 97.84 |
| *Bacillus velezensis* V417 | 3.90789 | 100 | 97.63 | 100 | 97.84 |
| *Bacillus velezensis* VCC-2003 | 3.92141 | 100 | 97.55 | 100 | 98.31 |
| *Bacillus velezensis* W1 | 4.23743 | 99 | 98.02 | 100 | 98.12 |

| | | | | | |
|---|---|---|---|---|---|
| *Bacillus velezensis* WRN014 | 4.06354 | 100 | 99.53 | 100 | 99.72 |
| *Bacillus velezensis* WS-8 | 3.92979 | 100 | 97.79 | 100 | 97.56 |
| *Bacillus velezensis* X030 | 3.95264 | 100 | 99.13 | 100 | 99.44 |
| *Bacillus velezensis* Y14 | 3.95716 | 100 | 99.61 | 100 | 99.06 |
| *Bacillus velezensis* Y2 | 4.23862 | 99 | 98.02 | 100 | 98.12 |
| *Bacillus velezensis* YAU B9601-Y2 | 4.24277 | 99 | 98.02 | 100 | 98.12 |
| *Bacillus velezensis* YJ11-1-4 | 4.00664 | 100 | 97.24 | 100 | 97.93 |
| *Bacillus velezensis* ZeaDK315Endobac16 | 3.92518 | 100 | 97.87 | 100 | 98.31 |
| *Bacillus velezensis* ZF2 | 3.92977 | 100 | 97.79 | 100 | 97.56 |
| *Bacillus velezensis* ZJU1 | 4.06415 | 99 | 98.02 | 100 | 98.12 |
| *Bacillus velezensis* ZL918 | 3.92271 | 100 | 99.37 | 100 | 99.06 |
| *Bacillus subtilis 168* | 4.21561 | 95 | 71.99 | 40 | 65.82 |
| *Bacillus licheniformis ATCC-14580* | 4.2226 | 46 | 66.88 | 63 | 67.46 |
| *Bacillus pumilus SAFR-032* | 3.70464 | / | / | / | / |
| *Bacillus cereus ATCC-14579* | 5.42708 | / | / | / | / |
| *Bacillus megaterium ATCC-14581* | 5.74664 | / | / | / | / |
